# *S. cerevisiae* serves as keystone species for spoilage resistance in experimental synthetic wine yeast communities

**DOI:** 10.1101/2024.07.04.602080

**Authors:** Alanna M. Leale, Eléonore Pourcelot, Stéphane Guezenec, Delphine Sicard, Thibault Nidelet

## Abstract

Species diversity is a commonly stated contributor to the fate of an invader, and thus community resistance, in both microbial and non-microbial communities. Termed the “diversity-invasion hypothesis”, a positive relationship between diversity and resistance to invasion is observed when an introduced species exhibits lower levels of survival in resident communities with higher species richness. The diversity-invasion hypothesis is an attractive perspective with convincing theory and examples, yet an “invasion paradox” of contrasting results means that a positive role of diversity against invasion is still not a certainty and under debate. In this study we investigated the relationship between resistance to invasion and resident community species richness versus species identity (i.e., keystone species). Using synthetic communities comprised of combinations of four wine yeasts (*Saccharomyces cerevisiae, Lachancea thermotolerans, Torulaspora delbrueckii, Starmerella bacillaris*), we tracked over 21 days the presence of introduced *Brettanomyces bruxellensis* spoilage yeast and *Lactiplantibacillus plantarum* lactic acid bacteria to ask the following: 1. Does yeast community species richness impact the establishment of *B. bruxellensis* yeast and *L. plantarum* bacteria during wine fermentation? 2. How does yeast species identity influence such establishment? We found that species identity rather than richness drove the prevention of establishment of *B. bruxellensis* and *L. plantarum*, with *S. cerevisiae* playing a critical keystone species role. Aside from spoilage prevention by *S. cerevisiae*, the four resident yeast species demonstrated a strict dominance ranking of competitive exclusion regardless of background community composition. Our research lends evidence against the commonly predicted positive relationship between species richness and resistance to invasion. Furthermore, as spontaneously fermented natural wines and diverse starter cultures gain popularity, our findings support a remaining importance of *S. cerevisiae* in preventing *B. bruxellensis* spoilage..

## INTRODUCTION

Ecological communities are collections of species or individuals that interact in a particular environment to contribute both individual and collective community functions. Communities may exist stably without changing species composition or function, or they may experience shifts in response to disturbances. The introduction and potential establishment of a novel species, or invader, is a common type of disturbance. Just as in macroecological communities, microbial communities also experience plentiful potential invasions, which involves the processes of introduction, establishment, spread, and impact by the introduced species (Mallon, Elsas and Salles 2015; Kinnunen *et al*. 2016; Vila *et al*. 2019). Moreover, species diversity is a commonly stated contributor to the fate of an invader, and thus resistance of both microbial (Jousset *et al*. 2011; van Elsas *et al*. 2012; Mallon, Elsas and Salles 2015; Kinnunen *et al*. 2016) and non-microbial communities (Kennedy *et al*. 2002; Tilman 2004; Petruzzella *et al*. 2020; Ernst *et al*. 2022). Termed the “diversity-invasion effect”, a positive relationship between diversity and resistance to invasion is observed when an introduced species exhibits lower levels of survival in resident communities with higher species richness (Kennedy *et al*. 2002; Mallon, Elsas and Salles 2015). Similarly, “biotic resistance theory” predicts that more phylogenetically diverse communities will be more resistant to invasion due to denser filled niche space, limiting colonisation by introduced species (Elton 1958; D’Antonio and Thomsen 2004; Petruzzella *et al*. 2020).

Resistance is considered the degree of change in a variable after versus before a perturbation (Allison and Martiny 2008; Donohue *et al*. 2016; Philippot, Griffiths and Langenheder 2021), thus in the context of invasion success, resistance is measured as the abundance of a novel species. As a community increases in species diversity, a wider range of resources are consumed, thereby limiting opportunities for a novel species to establish, and hence builds the basis of the diversity invasion effect and biotic resistance theory (Mallon, Elsas and Salles 2015; Petruzzella *et al*. 2020). Alternatively, overall diversity may play a negligible role due to influences of keystone species (Emery and Gross 2007; Ernst *et al*. 2023), the presence of a phylogenetically closely related species to the invader (Petruzzella *et al*. 2020), complex food webs (Thébault and Loreau 2005), resource heterogeneity (Jiang and Morin 2004; Fridley *et al*. 2007), and interaction strength (Mallon *et al*. 2015; Ratzke, Barrere and Gore 2020). The positive role of diversity against invasion is still not a certainty and under debate, probably because a combination of opposing factors. Although much insight has been gained using plant communities (Kennedy *et al*. 2002; van Ruijven, De Deyn and Berendse 2003; Petruzzella *et al*. 2020; Ernst *et al*. 2022), as well as diverse environmental microbial communities (van Elsas *et al*. 2012; Eisenhauer *et al*. 2013; Mallon *et al*. 2015; Xing *et al*. 2021), disentangling the multiple factors contributing to invasion resistance remains difficult in such complex systems. Therefore, defined synthetic microbial communities could help test single factors of invasion resistance factors at a time.

The composition of a resident community plays a critical role in its resistance to invasion. Community composition can entail several components, including the number or richness of species (i.e., How many?), the evenness of species (i.e., Dominant or rare types?), the phylogenetic diversity of species (i.e., How different?), the population density (i.e., How abundant?), and the identity of species (i.e., Who’s there?). We were specifically interested in exploring how a community’s resistance to invasion is influenced by identity effects of keystone species (Emery and Gross 2007; Ernst *et al*. 2023) compared to species richness. Similar to other recent research using synthetic microbial communities (Weiss *et al*. 2023), we *do not* use a restricted definition of a keystone species since we do not require a keystone species to have a low relative abundance compared to its influence on community function (Mills, Soulé and Doak 1993; Mouquet *et al*. 2013). We instead focus on microbial communities in which we consider a keystone species more generally as a species that, when absent, significantly alters community composition and/or function (Berry and Widder 2014; Weiss *et al*. 2023). In the context of microbial community invasion resistance, simply the presence versus absence of a keystone species can therefore be the influential factor determining whether an invader successfully establishes.

To explore the diversity-invasion resistance versus identity-invasion effect, we used a model synthetic community composed of wine yeasts. Wine yeast communities serve as a useful model system to test ecological questions due to their relatively manageable diversity, wealth of knowledge on their biochemistry or system function, and established synthetic media for laboratory experiments (Bagheri *et al*. 2020; Conacher *et al*. 2021; Pourcelot *et al*. 2023; Ruiz *et al*. 2023). It is already well known that starting extracted grape juice, termed grape must, initially harbours high diversity of yeast and bacteria, but this diversity drops as the fermentation progresses. Overtime, there is a rise and final dominance of *S. cerevisiae* as it withstands fermentation conditions of high alcohol concentration and low oxygen (Albergaria and Arneborg 2016; Conacher *et al*. 2021). Despite being at negligible or even undetectable levels in grape must (Fleet 2003; Zott *et al*. 2008; Drumonde-Neves *et al*. 2021), *S. cerevisiae* is considered necessary for complete fermentation “to dryness” (<2 g/L sugar) and is commonly added as a starter culture by wine producers (Albergaria and Arneborg 2016; Ciani *et al*. 2016; Binati *et al*. 2020). Furthermore, *S. cerevisiae* is known to prevent spoilage by achieving a quick, full fermentation which consumes available resources and leaves little opportunity for other species to proliferate (Ivey, Massel and Phister 2013; Williams, Liu and Fay 2015; Albergaria and Arneborg 2016). Additionally associated to *S. cerevisiae*’s strong fermentative capacities is its production of ethanol – an antimicrobial compound inhibiting growth of many other microbes (Du Toit and Pretorius 2000; Pereira, Freitas and Paschoalin 2021). The combination of strong resource competition and creation of a harsh environment with high ethanol concentrations by *S. cerevisiae*, together prevent spoilage. In ecological terms, the common problem of spoilage during wine production and during storage is an example of community instability against invasion. Thus, *S. cerevisiae* demonstrates a keystone species effect and helps make wine yeast communities a good model system to test diversity versus identity effects to invasion resistance.

Oenological fermentations are vulnerable to invasion by so-called spoilage species, such as the common problematic yeast *Brettanomyces bruxellensis*. To potentially combat spoilage and improve other product characteristics, more importance is being placed on promoting endemic yeast diversity and “non-*Saccharomyces”* yeasts in wine fermentations (Galati *et al*. 2019; Roudil *et al*. 2020). “Non- *Saccharomyces*” is a common term in wine science literature, but we avoid its use because the term is not ecologically based and actually typically refers specifically to the *S. cerevisiae* species, not the entire *Saccharomyces* genus (Jolly, Varela and Pretorius 2014). For the fundamental community ecology focus of our research, we here instead prefer referring to yeasts more specifically as non-*S. cerevisiae,* or communities being “*S. cerevisiae-*free”. Although naturally occurring non-*S. cerevisiae* yeasts can contribute favourably to aroma and sensory measures (Binati *et al*. 2020; Roudil *et al*. 2020), they can also pose problems with slow or stuck fermentations (Ciani, Beco and Comitini 2006; Medina *et al*. 2012; Taillandier *et al*. 2014). The dominance of *S. cerevisiae* has been shown to be influenced by ecological interactions with non-*S. cerevisiae* species (Boynton and Greig 2016; Bagheri *et al*. 2020; Conacher *et al*. 2021; Ruiz *et al*. 2023). If *S. cerevisiae* growth contributes to quick resource consumption and ethanol production, yet its growth also depends on its community, that brings to question - does species richness and consequent interactions also influence invasion by spoilage microorganisms?

In this work, we investigated the relationship between resistance to invasion and resident community species richness versus species identity (i.e., keystone species) using as model system synthetic wine yeast communities composed of four yeast species naturally found in grape must: *Saccharomyces cerevisiae*, *Lachancea thermotolerans*, *Starmerella bacillaris,* and *Torulaspora delbrueckii* (Pourcelot *et al*. 2023, 2024). The yeast *Brettanomyces bruxellensis* and the lactic acid bacteria *Lactiplantibacillus plantarum* were introduced as “invaders”. *S. cerevisiae* was a clear choice as an exemplary keystone species due to its strong fermentative performance and necessity in full fermentations. *L. thermotolerans* is a moderate fermenter and a popular non-*S. cerevisiae* yeast (Urbina, Calderón and Benito 2021). *S. bacillaris* is abundant in grape must (Csoma and Sipiczki 2008; Urso *et al*. 2008) and a weak-moderate fermenter, yet it was an interesting member to include due to its unique sugar preference for fructose rather than glucose metabolism (Englezos *et al*. 2018). Lastly, *T. delbrueckii* is a moderate fermenter that is phylogenetically and phenotypically (nitrogen source use) more similar to *S. cerevisiae* (Kemsawasd *et al*. 2015; Ramírez and Velázquez 2018). The high alcohol tolerance and ability to grow on limited resources of *B. bruxellensis* makes it a common spoilage organism in wine fermentation, including after bottling, where it produces undesirable “barnyard” or “sweaty-leather” aromas (Wedral, Shewfelt and Frank 2010; Malfeito-Ferreira and Silva 2019; Harrouard *et al*. 2023). *L. plantarum* was chosen due to its growing appeal for malolactic fermentation - a critical and desirable process in red wine fermentation (du Toit *et al*. 2011; Bravo-Ferrada *et al*. 2013; Urbina, Calderón and Benito 2021). Malolactic fermentation by lactic acid bacteria converts tart malic acid to the softer lactic acid and reduces the wine’s acidity (Lonvaud-Funel 1999; Virdis *et al*. 2021). However, a fine balance must be achieved because overgrowth of *L. plantarum* and other lactic acid bacteria also leads to undesirable levels of acetic acid (du Toit *et al*. 2011; Bartowsky, Costello and Chambers 2015; Urbina, Calderón and Benito 2021); interestingly then, it can be considered either a desired addition, or a spoilage invader. Furthermore, the *L. plantarum* bacteria was predicted to have less resource overlap with the resident yeast community, which may therefore contribute to its successful establishment or not. Using different communities varying for their species richness, we aimed to address two main questions: a) Does yeast community species richness impact the invasion of *B. bruxellensis* yeast and *L. plantarum* bacteria during wine fermentation? b) How does species identity (i.e., absence/presence) influence such invasion?

## METHODS

### Preparation of synthetic grape must (SGM) media

A synthetic grape must (SGM) media was used for all fermentations (prepared as in Bely et al. 1990) and adjusted to pH 3.3. Containing: 100 g/L glucose, 100 g/L fructose, 0.18g/L malic acid, 0.18g/L citric acid, and 425 mg/L of yeast assimilable nitrogen (mixture of ammonium chloride and amino acids). One large batch was prepared and kept frozen in smaller volumes until defrosted for each round of fermentations. Four separate rounds of fermentations were completed, thus providing four replicate data points. The day before inoculations, 250 mL of SGM was pasteurised in each fermenter, aerated for 20min, and stored overnight at 4°C. On the day of inoculations (d00), 850µL of β-sitosterol (final concentration of 5 mg/L) solution was added to each fermenter.

### Strains and fluorescent marking

Four yeast species comprised the “resident community” in our study: *Saccharomyces cerevisiae*, *Lachancea thermotolerans, Starmerella bacillaris,* and *Torulaspora delbrueckii* (details in Table 2). The strains used for *S. cerevisiae, L. thermotolerans, T. delbrueckii,* and *S. bacillaris* were each uniquely fluorescently tagged with an integrated fluorescent protein gene. This was done previously as in Pourcelot et al. 2023.

Two other microbes of interest were also included: the spoilage yeast *Brettanomyces bruxellensis* and the lactic acid bacteria *Lactiplantibacillus plantarum. L. plantarum* and *B. bruxellensis* strains were not fluorescently tagged. *L. plantarum* was chosen due to easier lab culturing compared to the better-known lactic acid bacteria *Oenococcus oeni*.

### Overnight cultures and day 0 (d00)

Overnight cultures of each species were prepared from a single colony (plates made less than a week prior from -80C° cell stock in YPD + 20% glycerol) in growth conditions described in Table 1. On day 00, the cell density of these overnight cultures was measured by flow cytometry. From the estimated cells/mL by cytometer, the correct volume of each species’ culture was added to a 50mL tube to reach a final 10^3^ cells/mL of *B. bruxellensis*, 10^3^ cells/mL of *L. plantarum*, and a 10^6^ cells/mL total of the remaining community members later in 250mL of SGM media (see Table 2). For example, in community C17 this was approximately 3.33 x 10^5^ *L. thermotolerans*, 3.33 x 10^5^ *S. bacillaris*, 3.33 x 10^5^ *T. delbrueckii* for a total of 10^6^ cells/mL. The mixed cultures were centrifuged (4500rpm, 5min, 4°C) then supernatant poured off, washed with 5mL of 9g/L NaCl, centrifuged again then supernatant poured off, and lastly 5mL of sterile SGM media added. The cells were resuspended in the SGM media by vortex and each community added to a separate pasteurised fermenter containing 250mL of SGM media. This created the d00 communities. An approximately 5mL sample was removed for d00 flow cytometry and plate counts. Bells were added to fermenters and their weights recorded in the Fermini computer software (internally developed), then placed at 28°C on 280 rpm spinners.

**Table 1:**
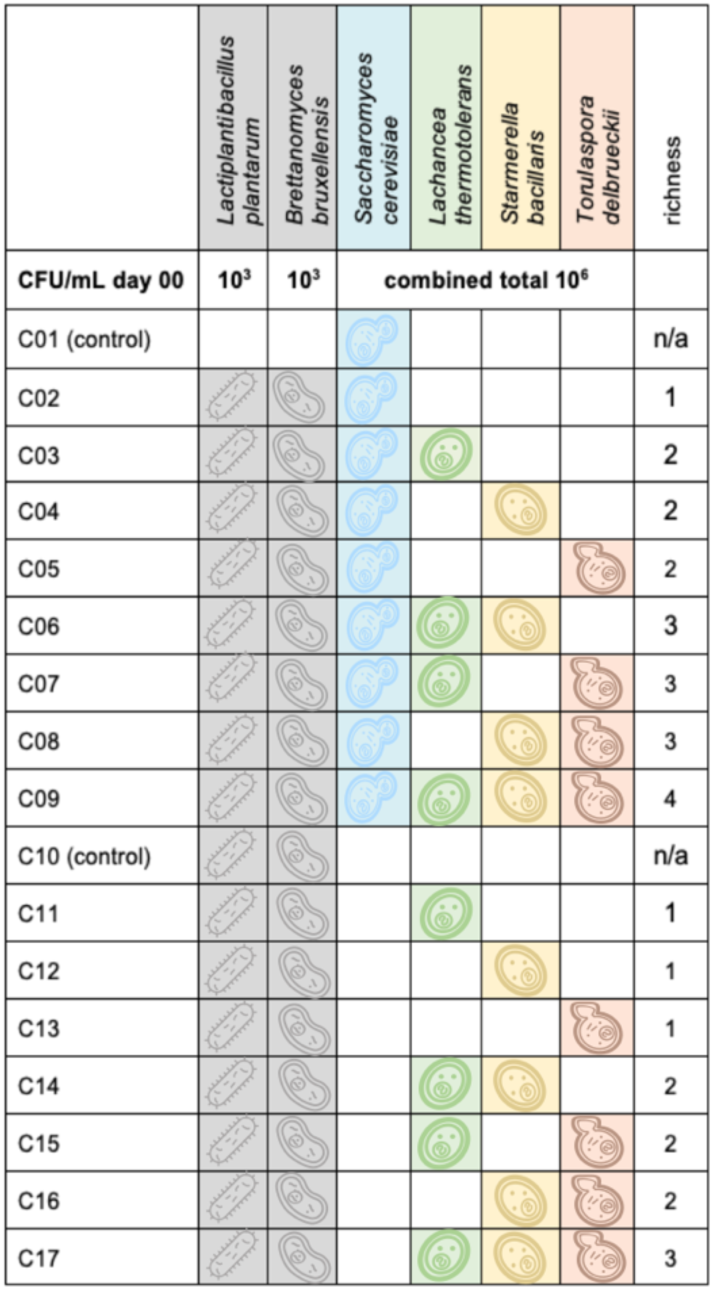
community compositions of 17 synthetic communities. Associated colours and icons for yeast species are used consistently across all figures.

**Table 2:**
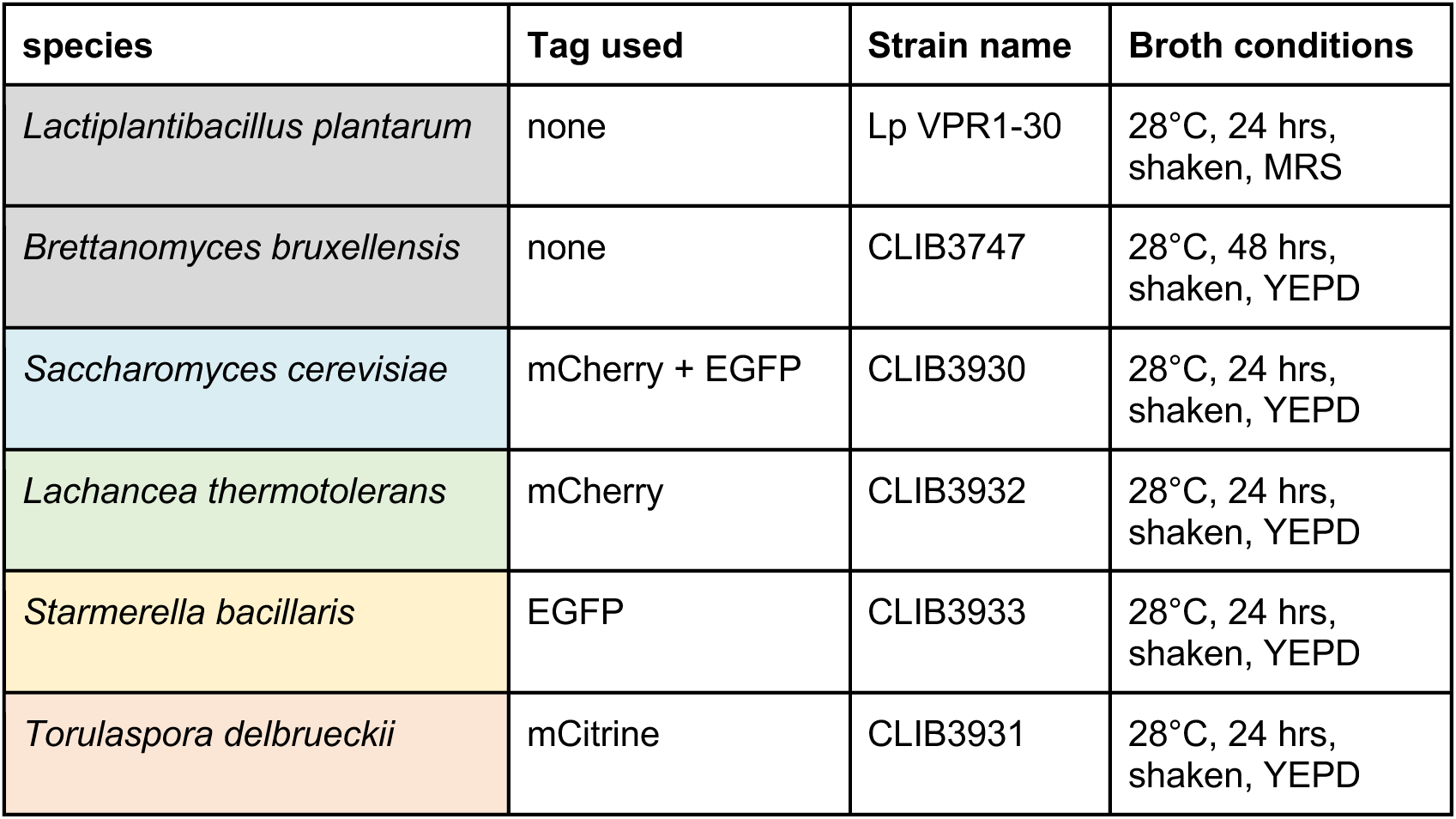
strains used and culture conditions.

| species | Tag used | Strain name | Broth conditions |
| --- | --- | --- | --- |
| *Lactiplantibacillus plantarum* | none | Lp VPR1-30 | 28°C, 24 hrs, shaken, MRS |
| *Brettanomyces bruxellensis* | none | CLIB3747 | 28°C, 48 hrs, shaken, YEPD |
| *Saccharomyces cerevisiae* | mCherry + EGFP | CLIB3930 | 28°C, 24 hrs, shaken, YEPD |
| *Lachancea thermotolerans* | mCherry | CLIB3932 | 28°C, 24 hrs, shaken, YEPD |
| *Starmerella bacillaris* | EGFP | CLIB3933 | 28°C, 24 hrs, shaken, YEPD |
| *Torulaspora delbrueckii* | mCitrine | CLIB3931 | 28°C, 24 hrs, shaken, YEPD |

### CO_2_ production/fermentation kinetics

At day d00, d01, d02, d07, d14, and d21, approximately 5mL was sampled from each fermenter for plate counts and/or flow cytometry. The weight was recorded before and after sampling in Fermini software. Estimated cumulative CO_2_ production (g/L) was calculated by the software at each time point and data exported to Excel.

### Plate counts

At day d00, d07, d14, and d21 the sample from each fermenter (I.e., community) was plated onto three selective medias to enumerate total yeast (excluding *B. bruxellensis*), *L. plantarum*, and *B. bruxellensis* (Table 3). A *B. bruxellensis* selective agar media was possible because of its resistance to cycloheximide and sensitivity of the other four yeast species (Morneau, Zuehlke and Edwards 2011). A conservative concentration of 50 µg/mL cycloheximide was previously confirmed from pilot experiments to stop growth of all yeast species except *B. bruxellensis*. The antibiotic chloramphenicol was also added to yeast selective medias to ensure full suppression of *L. plantarum* growth. Plates were counted on an automatic colony counter (Interscience Scan 300) and the calculated CFU values exported to Excel. A value of zero was only entered for empty plates at 10^0^ dilutions, otherwise the data point was considered missing.

**Table 3:**
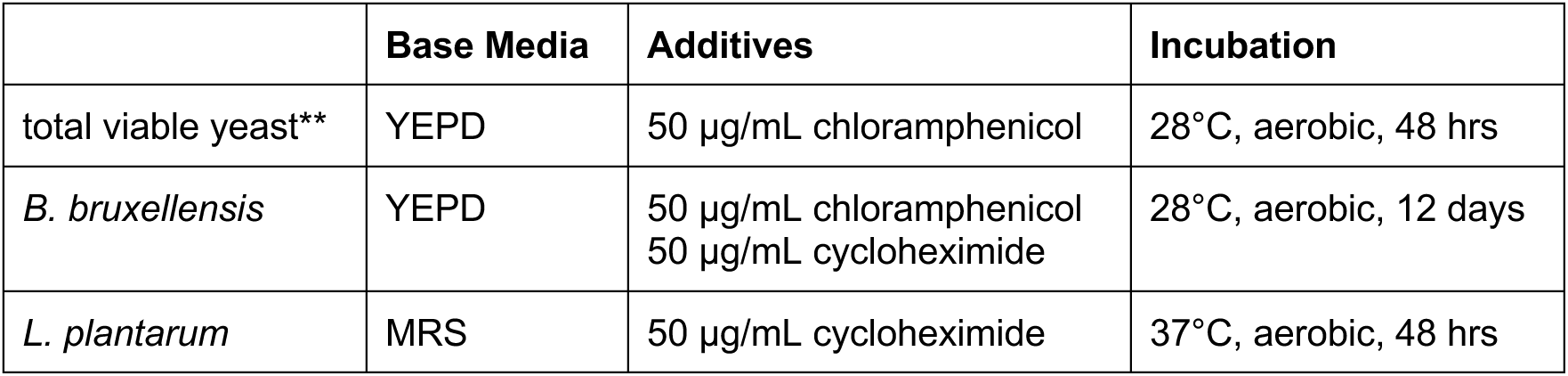
selective plating conditions. **Excludes *B. bruxellensis*

|  | Base Media | Additives | Incubation |
| --- | --- | --- | --- |
| total viable yeast** | YPD | 50 µg/mL chloramphenicol | 28°C, aerobic, 48 hrs |
| *B. bruxellensis* | YPD | 50 µg/mL chloramphenicol<br>50 µg/mL cycloheximide | 28°C, aerobic, 12 days |
| *L. plantarum* | MRS | 50 µg/mL cycloheximide | 37°C, aerobic, 48 hrs |

**Table 4:** Description of the set of channels and description of filters used for flow cytometry detection of the different yeast populations.

| Population | Fluorochrome | Channel | Voltage | Fluorescence |
| --- | --- | --- | --- | --- |
| Dead cells | PI | BL3 (488 – 695-40) | 340 | + |
| Live cells | PI | BL3 (488 – 695-40) | 340 | - |
| *S. cerevisiae* | mCherry + GFP | YL2 (600DLP-620/15) | 320 | + |
|  |  | BL1 (495DLP-530/30) | 260 | + |
| *L. thermotolerans* | mCherry | YL2 (600DLP-620/15) | 320 | + |
|  |  | BL1 (495DLP-530/30) | 260 | - |
| *S. bacillaris* | mCitrine | BL1 (495DLP-530/30) | 260 | + |
|  |  | VL2 (495DLP-512/25) | 400 | - |
| *T. delbrueckii* | EGFP | BL1 (495DLP-530/30) | 260 | + |
|  |  | VL2 (495DLP-512/25) | 400 | - |

As planned, the negative control community C10, containing only *B. bruxellensis and L. plantarum* exhibited no viable yeast colonies at day 00 (Fig. S1). Our negative control community C01 of *S. cerevisiae* alone, without *B. bruxellensis*, showed no growth throughout the experiment, lending confirmation that cycloheximide was selective for *B. bruxellensis* (Fig. S2). Our negative control community C01 of *S. cerevisiae* alone, without *L. plantarum*, showed no growth throughout the experiment, lending confirmation that our chloramphenicol media was selective for *L. plantarum*. (Fig. S3).

To note, we struggled finding correct dilution factors, leading to some missing data or lawns with closest CFUs estimated. Most notably, we believe a human error in Round 1 showed zero viable cells at day 21, which inaccurately guided our dilution factor choices at later rounds and resulted in lawns that were uncountable. Lawns were conservatively entered as the minimum possible colonies (i.e., 100) and the dilution factor (i.e., a lawn at 10^-1^ = 10^4^ CFU/mL). Also to note, we assume that there was early contamination by *S. cerevisiae* of the C10 community “negative” control (containing only slow growing *B. bruxellensi*s and the poor growing bacteria *L. plantarum)* in replicate round 3 and 4 (Fig. S1). The contamination is not surprising at all since *S. cerevisiae* is ubiquitous in the lab environment and is the strongest competitor in our synthetic must environment.

### Flow cytometry population dynamics

The fermenting must was sampled aseptically at day d00, d01, d02, d07, d14, d21 to determine number cells of each species with flow cytometry. Population cell number was monitored by flow cytometry with the Attune NxT™ Thermofisher® Flow Cytometer (Life Technologies, Singapore) equipped with an AttuneNXT Autosampler. Briefly, cells were washed in PBS (130 mM NaCl, 2.6 mM KCl, 7 mM Na2HPO4, 1.2 mM KH2PO4, pH 7.4; Sigma) and diluted to obtain cell concentrations about 1 to 5.10^5^ cells/mL. Numeration of live cells (i.e., viability) was assessed after staining cells with 1 μg/mL propidium iodide (PI, stored at 4°C protected from light; Calbiochem). Each population was previously tagged with one or two fluorescent proteins (Pourcelot et al. 2023) and then detected with a specific set of channels described in Table X. Gating was done with the AttuneNxT software.

Our gatings proved to be appropriate, since unexpected species were not found in corresponding communities (i.e., only *L. thermotolerans* was found in C11 where only *L. thermotolerans* was inoculated). There is probable underestimation of *T. delbrueckii* cell counts by flow cytometry due to its flocculating nature. We put our best efforts to thoroughly resuspend those *T. delbrueckii* containing samples by vortex and manual pipetting.

### Statistical analyses

All analyses were performed in R version 4.3.2 (R Core Team 2023). Significance was defined for all analyses as p-value < 0.05. Results outputs and more details of statistical analysis are found in Supplementary Material.

### Cumulative CO_2_

Separately for data points at day 2 and then day 21, a linear model testing the effect by community richness and each species’ presence on cumulative CO_2_ was fitted using the “lm” function from “lme4” R package. Cumulative CO_2_ was the continuous response variable with absence versus presence of each species (α, β, δ, ε) and richness (θ) as additive categorical fixed effects (Bates *et al*. 2015). γ = α + β + δ + ε + θ, where γ = cumulative CO_2_, α = presence of *S. cerevisiae* (0, 1), β = presence of *L. thermotolerans* (0, 1), δ = presence of T. *delbrueckii* (0, 1), ε = presence of *S. bacillaris* (0, 1), θ = richness level (1, 2, 3, 4).

A PERMANOVA was performed to evaluate the variation in cumulative CO_2_ of *S. cerevisiae*-free versus *S. cerevisiae*-containing communities. The “vegdist” function was used to calculate Euclidean distance of variation in cumulative CO_2_, followed by the “adonis” function from the “vegan” R package, with “bray" method. Presence of *S. cerevisiae* was the categorical effect variable (0, 1) with Euclidean distance as the continuous response variable (Oksanen *et al*. 2022).

### Total viable yeast, B. bruxellensis, L. plantarum plate counts

Inexactness of cell counts across time points (i.e., agar plate lawns, empty plates) led us to perform all statistical tests not as quantitative cell counts but instead as binary data (absence versus presence, presence being any colony growth detected). Fisher’s Exact Tests were completed using the “fisher.test” function of base R, to evaluate the effect of community richness and each species’ presence on the detection of viable yeast, *B. bruxellensis,* and *L. plantarum*. The Fisher’s exact test is recommended for smaller data sets below 1000 points, which is suitable for our study. A separate Fisher’s Exact test was completed separately for each output, day, and variable combination. A Bonferroni correction of the p-value significance threshold was performed to account for multiple tests (i.e., adjusted p-value threshold = 0.05 / # tests).

### Flow cytometry maximum total CFU/mL

The same as for *Cumulative CO_2_,* a linear model testing the effect on maximum CFU per mL of each species’ presence and community richness was fitted using the “lm” function from “lme4” R package. Maximum CFU reached was the continuous response variable with absence versus presence of each species and richness as additive categorical fixed effects (Bates *et al*. 2015).

## RESULTS & DISCUSSION

### S. cerevisiae-free communities ferment slower and more variably

In this study, fermentations with combinations of four resident wine yeast species (*S. cerevisiae*, *L. thermotolerans*, *S. bacillaris, T. delbrueckii*) were performed to investigate the impact of species richness and community composition on the invasion of two potential spoilage micro-organisms (*B. bruxellensis*, *L. plantarum)*. The communities varied in resident species richness (1, 2, 3, or 4 species) and composition, where for example, half the communities included *S. cerevisiae* (*S. cerevisiae*- containing) and the other half did not (*S. cerevisiae*-free) (community compositions listed in Table 1). Firstly, the fermentation performance of the 15 communities in synthetic grape must was tracked over 21 days through weight measurements that is proportional to CO_2_ production and calculated cumulative CO_2_ (Fig. 2). A higher rate of fermentation was observed in communities containing *S. cerevisiae* (C02- C09, Fig. 2 dashed lines). As expected, *S. cerevisiae*-free communities in comparison showed overall slower fermentation with a significant lower cumulative CO_2_ production at day 2 (C11-C17, Fig. 2 solid lines; F_6, 52_ = 82.79, p-value <2e-16, Table S4) and most did not reach comparable cumulative CO_2_ levels (i.e., fermentation level) as those containing *S. cerevisiae,* even by day 21. However, we saw that some *S. cerevisiae*-free communities (C13, C15, C16, C17) reached cumulative CO_2_ levels comparable to *S. cerevisiae*-containing communities at day 21 (using the cumulative CO_2_ plateau of C02-C09 as a reference). The common component across these four communities was the inclusion of *T. delbrueckii* in their starting compositions. They reached in average at day 21 a CO_2_ production of 95.40±6.45 g/L compared to 90.76±9.10 g/L for the communities without *T. delbrueckii* (F_6, 53_ = 4.75, p-value = 0.03251, Table S5). Additionally, *S. cerevisiae-*free communities (C11-C17) had greater variation compared to when *S. cerevisiae* was present (C02-C09) (p-value = 0.002, Table S6). The minimal variation in fermentation across *S. cerevisiae*-containing communities reflects the known standardisation effect that *S. cerevisiae* starter cultures have on wine fermentation (Ciani *et al*. 2010). When analysing across all timepoints, *S. cerevisiae* was the only species whose presence had a significant effect on cumulative CO_2_ (F_7, 352_ = 92.98, p-value = 3.88e-07, Table S7).

**Figure 1:**
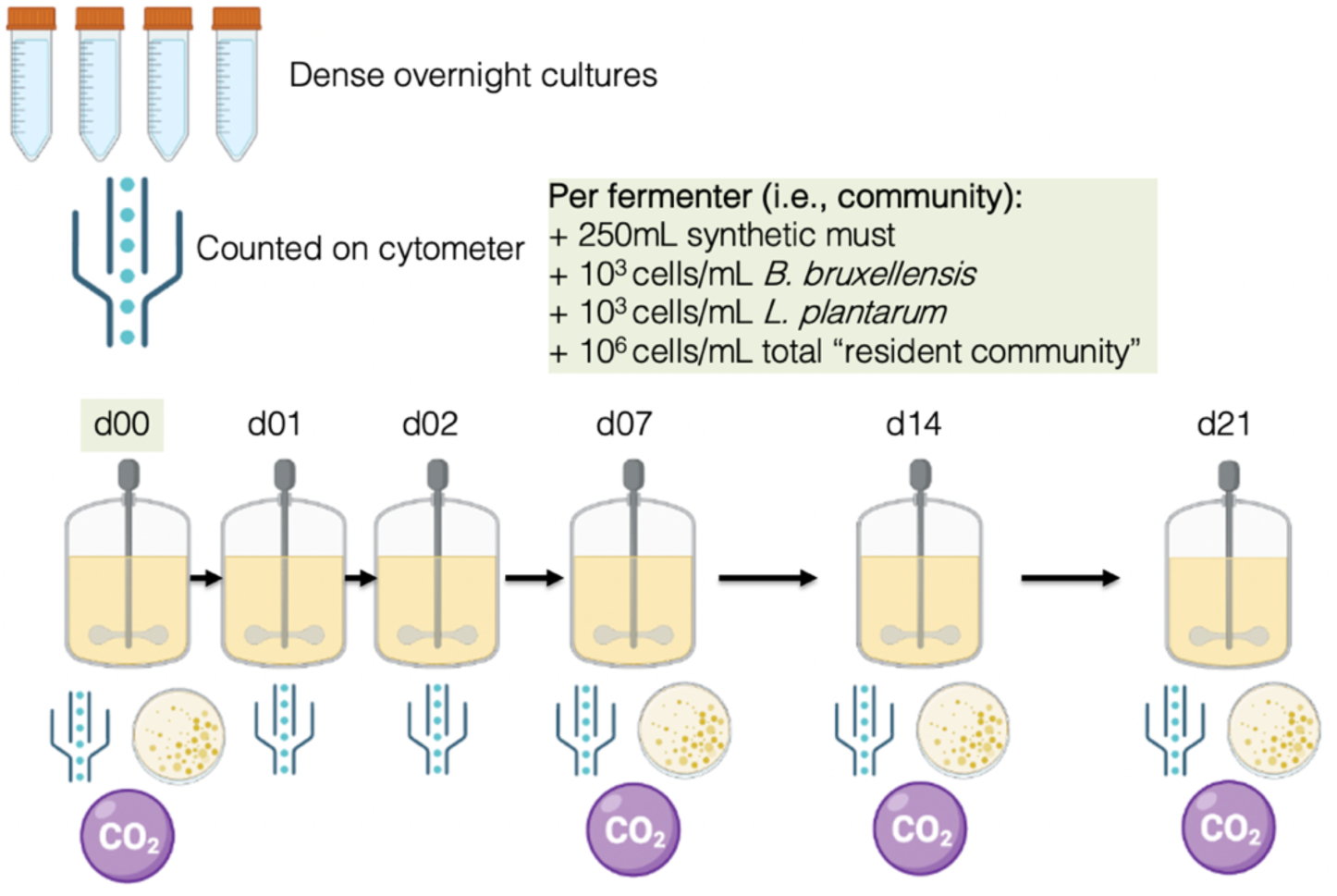
experimental setup. Figure created with BioRender.com.

**Figure 2:**
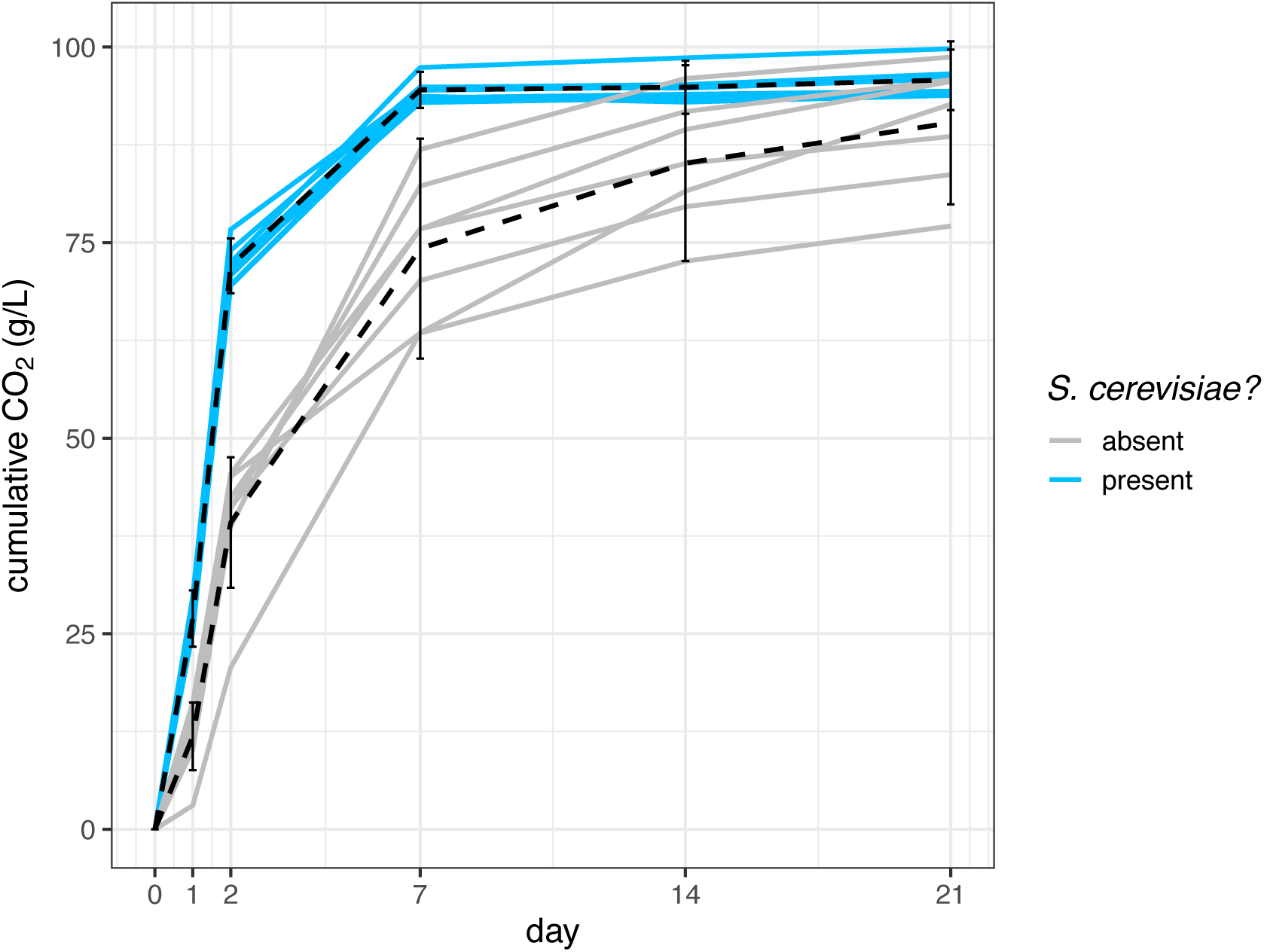
*fermentation is stunted in the absence of S. cerevisiae.* Cumulative CO2 produced over time demonstrates fermentation progression in all 17 synthetic communities. Solid lines are the mean of four replicates for each community. Dotted lines show mean across all *S. cerevisiae*-free (grey) or *S. cerevisiae*-containing (blue) communities, with error bars showing one standard deviation.

A recent large-scale study found that *S. cerevisiae* growth and consequent final sugar consumption was reduced when co-cultured with other yeast species (Ruiz *et al*. 2023). They suggested that the reduction in *S. cerevisiae* growth was due to heightened negative species interactions resulting from greater species richness. While we did not directly measure consumed sugars, we do still see in our cumulative CO_2_ data that all *S. cerevisiae*-containing communities reach fermentation plateau by day 7, regardless of community richness (F _1, 30_ = 0.0053, richness level p-values > 0.05, Table S8). The discrepancy may be because some strains used by Ruiz et al. 2023 were more strongly predicted to have negative interactions on *S. cerevisiae* (mainly *P. kudriavzevii, H. opuntiae,* and *M. pulcherrima),* whereas these species were not included in our study. Another possibility is that slight composition differences in their synthetic grape must media compared to ours (i.e., yeast assimilable nitrogen, trace elements, vitamins) influences competition dynamics since species interactions depend on the environment and available substrates. Overall, fermentation in *S. cerevisiae*-containing communities plateaued earlier and progressed more consistently than *S. cerevisiae-*free communities, which implies slower, more variable, and incomplete resource consumption across *S. cerevisiae*-free communities.

### Viable yeasts persist after 7 days in S. cerevisiae-free communities

Using colony counts on YEPD media agar plates, we estimated total viable yeast (excluding *B. bruxellensis*) in the 15 resident communities at day 00, day 07, day 14, and day 21 (i.e., total combined *S. cerevisiae, S. bacillaris, L. thermotolerans,* and *T. delbrueckii*) (Fig. 3). We confirmed that we successfully inoculated approximately 10^6^ cells/mL total resident community yeast cells at day 00 (Fig. S1). To assess the growth of resident yeasts over time, we scored replicate plates as presence (calculated CFU/mL > 0) or absence (calculated CFU/mL = 0) of yeast and calculated the frequency of replicates where viable yeast was detected for each community and time point (Fig. 3A). Viable yeast cells were not detected by day 21 in communities containing *S. cerevisiae* (C02-C09), which aligns well with established knowledge of wine fermentation dynamics demonstrating rapid growth and consumption of resources by *S. cerevisiae* (Albergaria and Arneborg 2016). Comparatively, in *S. cerevisiae*-free communities (C11-C17), viable yeast cells were still observed at day 14 and even day 21 (non-empty plate proportion: day 14 = 0.75, day 21 = 0.50), which we interpret as a slower consumption of resources and delayed approach to carrying capacity. Furthermore, our detection of total viable yeast cells supports the rapid fermentation by *S. cerevisiae* containing communities observed in CO_2_ fermentation performance data (Fig. 2). Early and fast consumption of resources by the resident yeast community is predicted to strengthen resistance against *B. bruxellensis* and *L. plantarum* because fewer remaining resources restricts opportunities for spoilage microorganisms to establish. Quick fermentation by *S. cerevisiae* will also contribute to high levels of ethanol earlier; however, this is likely not a main factor preventing *B. bruxellensis* growth since the spoilage yeast exhibits ethanol tolerances as strong as *S. cerevisiae* (Renouf *et al*. 2006; Childs, Bohlscheid and Edwards 2015; Smith and Divol 2016). Although strain dependent, ethanol stress tolerance can also be relatively high for *L. plantarum* or other lactic acid bacteria, with growth observed at ethanol concentrations of 10 to 13% (G-Alegría *et al*. 2004; Ngwenya, Nkambule and Kidane 2023).

**Figure 3:**
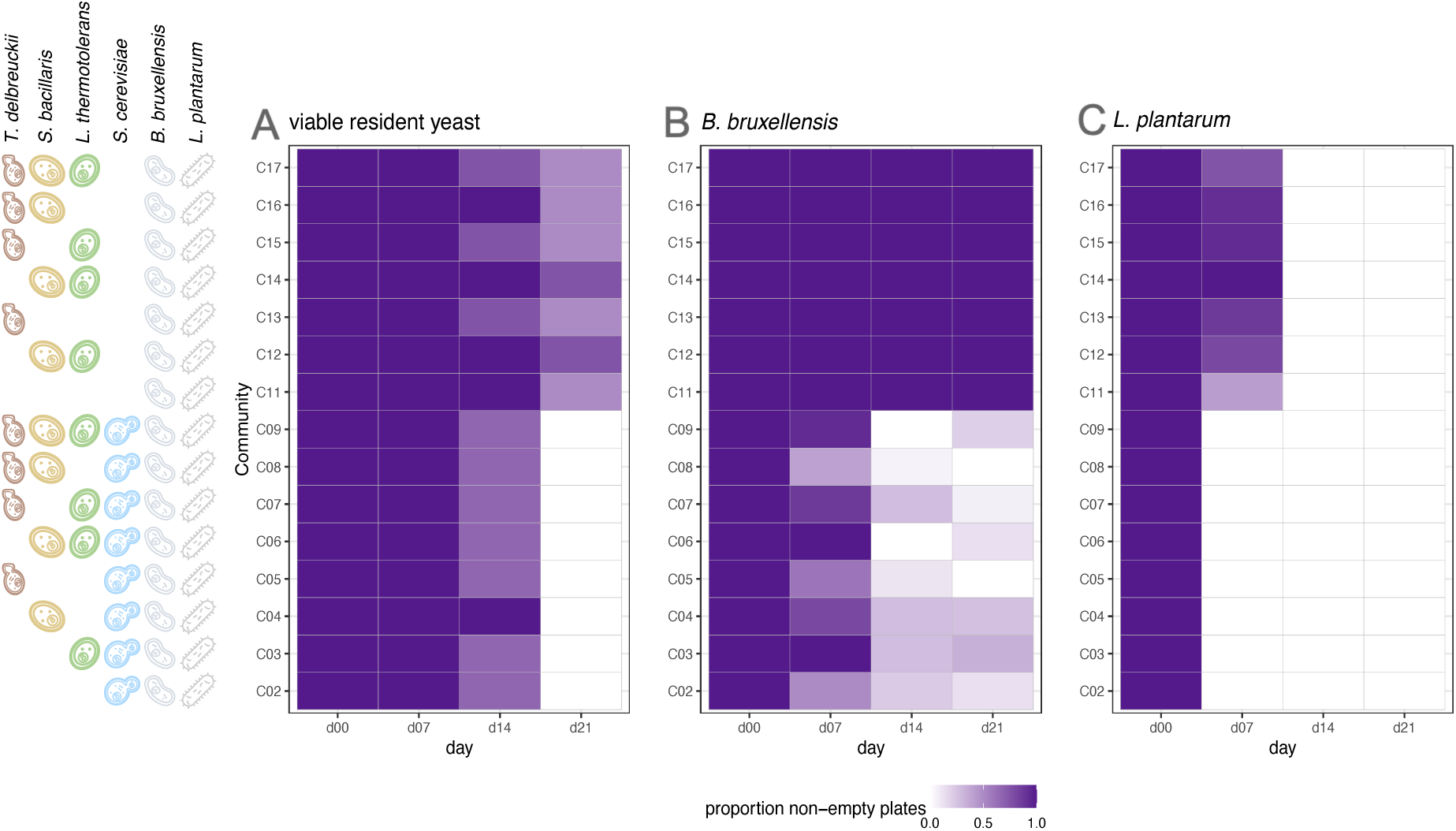
**A)** *viable yeast remain in S. cerevisiae-free communities.* Detected presence of total viable yeast (excluding *B. bruxellensis*) over time from YEPD agar plating and counting after 48 hours. **B)** *B. bruxellensis establishes in the absence of S. cerevisiae.* Detected presence of *B. bruxellensis* over time from selective agar plating on YEPD supplemented with cycloheximide and counted after approximately 14 days. **C)** elimination of *L. plantarum* is delayed in the absence of *S. cerevisiae.* Detected presence of *L. plantarum* over time from selective agar plating on MRS supplemented with chloramphenicol and counted after 48 hours. Species compositions of each community indicated as icons on left side.

The presence of a viable resident yeast community was influenced by a negative identity effect of *S. cerevisiae* presence at days 14 and 21 (Fisher’s Exact Tests: Table S1). Species richness of the community had a negative effect of viable yeast of the resident community at day 21 (Fisher’s Exact Tests: Table S1). However, when only *S. cerevisiae*-free communities and only *S. cerevisiae*-containing communities were separately analysed, the negative effect on viable yeast by species richness disappeared (Fisher’s Exact Tests: Tables S2 and S3), lending evidence that richness effects are due to a confounded greater probability of *S. cerevisiae* inclusion, i.e., identity effect, rather than richness alone. Having observed that viable yeast remained in *S. cerevisiae*-free communities until day 21, we next looked at the presence of *B. bruxellensis* overtime across our 15 communities of varied species richness and composition.

### B. bruxellensis proliferates in S. cerevisiae-free communities

We then estimated the abundance of the spoilage yeast *Brettanomyces bruxellensis* overtime. The analysis of *B. bruxellensis* population density at day 00 showed that each community was successfully inoculated with approximately 10^3^ cells/mL of *B. bruxellensis* (Fig. S2). As for total viable yeast, we then calculated the frequency of replicates where *B. bruxellensis* was detected as present (calculated CFU/mL > 0) (Fig. 3B). After seven days in communities containing *S. cerevisiae* (C02-C09), *B. bruxellensis* was already starting to decrease (Fig. 3B). The gradual elimination of *B. bruxellensis* in *S. cerevisiae*-containing communities continued further by day 14 and day 21 (non-empty plate proportion: day 07 = 0.40 to 1.00, day 14 = 0.00 to 0.28, day 21 = 0.00 to 0.33). In comparison, *B. bruxellensis* populations persisted in *S. cerevisiae*-free communities until day 21 (minimum non-empty plate proportion = 1.00). From day 7 onwards, *B. bruxellensis* demonstrated cell counts remaining above target inoculation values (Fig. S2). There was a significant negative effect by species richness and the presence of *S. cerevisiae* on *B. bruxellensis* detection at days 07, 14, and 21 (Fisher’s Exact Tests: Table S1). Yet, the negative effect by species richness on *B. bruxellensis* disappeared when *S. cerevisiae*-free and *S. cerevisiae*-containing communities were separately analysed (Fisher’s Exact Tests: Tables S2 and S3). Greater species richness is confounded with a greater probability of *S. cerevisiae* being present, thus our results support a strong keystone species role of *S. cerevisiae* that causes an overall observed effect of species richness. The persistence or even growth of *B. bruxellensis* in *S. cerevisiae*-free communities aligns with the concept of breached biotic resistance of the resident community and successful establishment of the invader (Mallon, Elsas and Salles 2015). Entering the growth and spread phase exhibits *B. bruxellensis’* ability to access local resources in communities lacking *S. cerevisiae*, which aligns with our results of fermentation progression (Fig. 2) and presence of viable yeast in the resident community (Fig. 3A).

Our results suggest a critical role for *S. cerevisiae* in the prevention of spoilage by *B. bruxellensis* in wine fermentation, however we must consider limitations from the simplicity of our synthetic grape must media (SGM). In real wine production settings, *B. bruxellensis* is still a problematic spoilage organism, even if *S. cerevisiae* is present (Wedral, Shewfelt and Frank 2010; Malfeito-Ferreira and Silva 2019; Harrouard *et al*. 2023). We cannot perfectly mimic the high resource complexity found in natural grape must from vineyards, which may provide opportunities for *B. bruxellensis* to persist since the species is known to have minimal nutrient requirements (Childs, Bohlscheid and Edwards 2015; Smith and Divol 2016). None-the-less, as enthusiasm rises for natural wine fermentation and the beneficial roles of naturally occurring non-*S. cerevisiae* yeasts for wine characteristics (Galati *et al*. 2019; Roudil *et al*. 2020), wine producers should not underestimate the importance of *S. cerevisiae.* Creative strategies to permit the contributions of both *S. cerevisiae* and other present yeast species are being sought, such as delayed inoculation of starter cultures containing *S. cerevisiae* (Taillandier *et al*. 2014; Conacher *et al*. 2021), reduced temperatures (Alonso-del-Real *et al*. 2017), or oxygenation early in fermentation (Shekhawat, Bauer and Setati 2017) to promote growth of less competitive species (Englezos *et al*. 2022). In addition to *S. cerevisiae*, other overlooked *Saccharomyces* species with varying fermentation capabilities should be studied for their potential role in spoilage prevention. Our model wine yeast community model system could replace *S. cerevisiae* with various alternative high or low fermenter *Saccharomyces* species and see if keystone species effects are also observed generally for the *Saccharomyces* genus, or if they are specific to *S. cerevisiae.* Such an approach would enable testing theories of predicted niche overlap as a function of phylogenetic distance (Elton 1958; D’Antonio and Thomsen 2004; Petruzzella *et al*. 2020) versus phenotypic metabolic traits (i.e., sugar consumption efficiency, alcohol tolerance, nitrogen availability limitations).

### L. plantarum is eliminated slower in S. cerevisiae-free communities

We also explored the effect of species richness versus species identity on the persistence of lactic acid bacteria during wine fermentation by tracking *Lactiplantibacillus plantarum* in our synthetic wine yeast communities. We estimated abundances of *L. plantarum* overtime using colony counts on agar media selective for lactic acid bacteria, and then calculated the frequency of replicates where *L. plantarum* was detected as present (calculated CFU/mL > 0) (Fig. 3). We first confirmed that the initial *L. plantarum* population was successfully inoculated at approximately 10^3^ cells/mL (Fig. S3). *L. plantarum* was rapidly eliminated by day 7 from communities containing *S. cerevisiae* (C02 to C09) with a significant negative effect of *S. cerevisiae* and species richness on the presence of *L. plantarum* (Fisher’s Exact Tests: Table S1). In *S. cerevisiae*-free communities (C11-C17), *L. plantarum* abundances remained detected (Fig. 3C) (minimum non-empty plate proportion = 0.42) but mostly below target starting levels (Fig. S3), thus it did not proliferate in these communities. By day 14, *L. plantarum* was no longer detected in any of the fifteen communities, regardless of resident community composition. Establishment of an invader progresses either because it possesses unique metabolic traits that permit it fill an empty niche, or more commonly, because it outcompetes at least one member of the resident community (Kinnunen *et al*. 2016). We had speculated that unique bacterial metabolism of *L. plantarum*, compared to resident yeast species, could potentially enable it to fill an unutilised niche. Contrary to our expectation, it does not appear that *L. plantarum* was able to breach biotic resistance of the resident community. We interpret its inability to enter a growth phase to demonstrate that local resources were not available to the bacteria and that the resident yeasts, regardless of species identity and richness, eventually outcompete *L. plantarum*. We did not measure gas or liquid phase metabolic profiles in this study, but it would be useful for future studies to assess if there are metabolic traces of *L. plantarum* in communities where it persisted to day 7. The aromatic and flavour contributions of *L. plantarum* is worthy of investigation since lactic acid bacteria can produce both desirable and undesirable characteristics to wine. Overgrowth of lactic acid bacteria can produce excess acetic acid (du Toit *et al*. 2011; Bartowsky, Costello and Chambers 2015; Urbina, Calderón and Benito 2021), yet malolactic fermentation is also essential to reduce the acidity of wine by converting malic acid to lactic acid (Lonvaud-Funel 1999; Virdis *et al*. 2021). An interest for diverse lactic acid bacteria starters is forming since typically just *Oenococcus oeni* is used (Bartowsky, Costello and Chambers 2015; Cappello *et al*. 2017), thus it is of value to find other suitable species and strains for starter culture communities. Finding suitable strains requires experiments that also measure organic acids (malic acid, lactic acid, acetic acid, etc.) and pH to assess the effectiveness of various lactic acid bacteria (including *O. oeni* and *L. plantarum*) for converting malic acid, versus their production of undesirable compounds. *L. plantarum* is often isolated together with *S. cerevisiae* in various fermented foods (Liu *et al*. 2022), yet our results suggest that *L. plantarum* is not a strong competitor with our synthetic wine yeast communities and must environment, especially when *S. cerevisiae* is present. Contrasting results between our synthetic experiment versus natural wine fermentation could be an outcome of the simplicity of our SGM media and our inoculation of *S. cerevisiae* at higher ratios than typically found in natural grape must (Albergaria and Arneborg 2016; Conacher *et al*. 2021; Drumonde-Neves *et al*. 2021).

### Hierarchical dominance across yeast species observed by flow cytometry

Our four resident species were each uniquely fluorescently marked, enabling us to estimate their abundances over time using flow cytometry. As expected, flow cytometry data demonstrated a quick dominance of *S. cerevisiae* and a nearly complete exclusion of the other yeast species (Fig. 4, blue lines, C02-C09). The only slight exception of exclusion by *S. cerevisiae* was in communities where *L. thermotolerans* briefly coexisted from day 01 to day 03, albeit at low abundance (C03, C06, C07, C09 - green lines). *T. delbrueckii* (red lines) and *S. bacillaris* (gold lines) were entirely outcompeted by *S. cerevisiae* within the first days of fermentation. The stronger competitive ability of *L. thermotolerans* over *T. delbrueckii* and *S. bacillaris* is further demonstrated by its dominance in *S. cerevisiae*-free communities C14, C15, and C17. *S. bacillaris* is evidently the poorest competitor, as it only reached detectable levels when in monoculture (community C12) and was outcompeted when paired with *T. delbrueckii* (community C16). Furthermore, in monocultures, *L. thermotolerans* reached the greatest maximum population size after *S. cerevisiae* (Fig. S5). The dominance of a species was not affected by the presence of others in the community and followed the ranking order from strongest to weakest competitor of: *S. cerevisiae, L. thermotolerans, T. delbrueckii, S. bacillaris*. Importantly, this dominance ranking held regardless of the community composition; for example, *L. thermotolerans* always outcompeted *T. delbrueckii,* regardless of which other yeast species were present.

**Figure 4:**
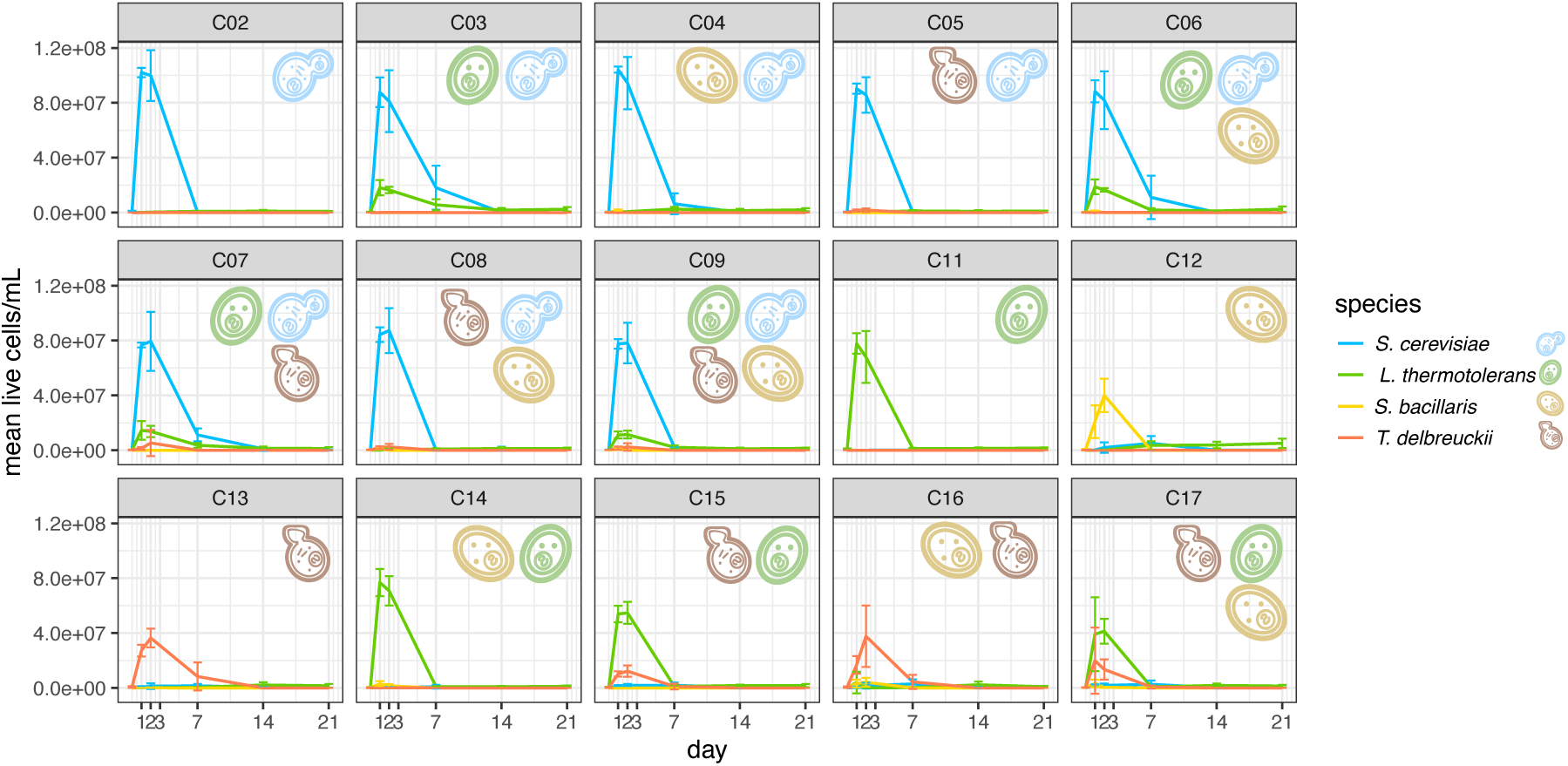
*Flow cytometry reveals species dominance ranking: S. cerevisiae > L. thermotolerans > T. delbrueckii > S. bacillaris.* Fluorescent flow cytometry estimates of resident community yeast species overtime. The mean live cells/mL of four replicate rounds of fermentation is plotted, with error bars showing one standard deviation. Icons indicate species composition of each community.

It was not surprising that *S. cerevisiae* quickly outcompeted the other species, given its strong fermentative ability and high inoculation concentration (equivalent to fellow resident species) compared to in natural grape must where it is a minority (Albergaria and Arneborg 2016; Conacher *et al*. 2021; Drumonde-Neves *et al*. 2021). We can compare our dominance rankings of the other three species to another study of pairwise interactions of *L. thermotolerans, S. bacillaris,* and *S. cerevisiae* (as well as other species); they similarly saw that *L. thermotolerans* could persist well through fermentation, whether inoculated or naturally present as indigenous strains (Bagheri *et al*. 2020). Our results demonstrated *S. bacillaris* as the poorest competitor even when in monoculture, whereas their research found that it could dominate if *S. cerevisiae* was absent. Our results additionally differ slightly from their finding that population dynamics between wine yeast species change in *S. cerevisiae-*free communities (Bagheri *et al*. 2020); we observed the same dominance of *L. thermotolerans > T. delbrueckii > S. bacillaris* when *S. cerevisiae* was present or not. Bagheri et al. 2020 used more complex and diverse natural communities, whereas our study’s simplicity of a four species synthetic community maybe misses complex ecological interactions that influence wine yeast species dynamics. Most importantly, our results align with a closely related study to ours that studied pairs of the same fluorescently marked strains; although they did detect interactions between the species in co-culture, there were consistent dominance rankings (Pourcelot *et al*. 2023).

Interestingly, despite *L. thermotolerans* reaching considerable maximum population sizes in communities C11 and C14 (Fig. 4, Fig. 5A), *B. bruxellensis* still proliferated (Fig. 3B). However, the maximum number of live cells in C11 and C13 is still below that of *S. cerevisiae*-containing communities (Fig. 5A). The persistence of *B. bruxellensis* indicates that resources were not exhausted, or at least could not be used quick enough, by the resident community. We therefore hypothesise that compared to *S. cerevisiae, L. thermotolerans* is less efficient in consumption of carbon and nitrogen sources, consequently leaving a greater opportunity for *B. bruxellensis* to establish. In other words, we could interpret that the resource niche overlap appears greater between *B. bruxellensis* and *S. cerevisiae,* compared to *B. bruxellensis* and *L. thermotolerans*. Resource use efficiency could be tested by growing *B. bruxellensis* in spent media of *L. thermotolerans* versus *S. cerevisiae* at multiple time points of fermentation. Successful growth of the spoilage yeast in spent media could reveal its window of opportunity for establishment and demonstrate niche overlap with alternative resident yeast species.

**Figure 5:**
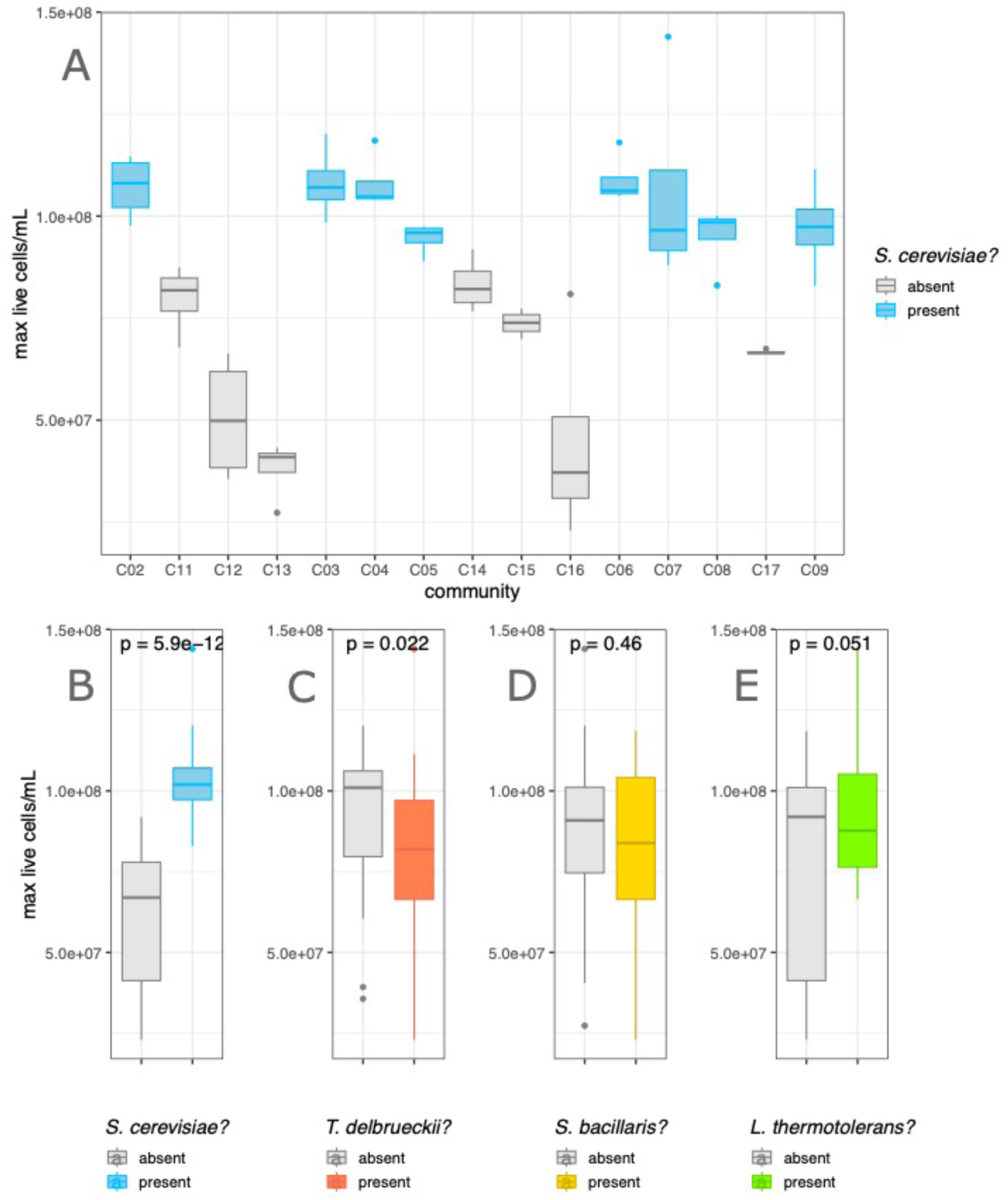
*Maximum CFU is increased only by the presence of S. cerevisiae*. Fluorescent flow cytometry estimates of maximum live cells/mL reached during fermentation of the resident community yeasts (i.e., excludes *B. bruxellensis*). Replication = 4 fermentation rounds per community. **A)** each community separately, ordered on the X-axis by community species richness (richness 1 = C02, C11, C12, C13; richness 2 = C03, C04, C05, C14, C15, C16; richness 3 = C06, C07, C08, C17; richness 4 = C09. **B) to E)** Data points pooled together based on absence versus presence of each resident yeast species. P-value is of t-test comparing the two groups.

To also note is the discrepancy in our data between estimated alive cells from the cytometer versus plate counts for total yeast. When observing cytometer readings on a log10 scale, it is revealed that all communities have up to 10^6^ cells/mL at day 14 and day 21 (Fig. S4), yet total viable yeast plate counts for communities C02-C09 show no live cells (Fig. 3A). The proliferation of *B. bruxellensis* in *S. cerevisiae*-free communities cannot explain the differences since both cytometer and viable yeast plate count measurements exclude the spoilage species; *B. bruxellensis* was not fluorescently tagged for cytometer detection, and yeast plate counts were performed at 48 hours, long before the growth of *B. bruxellensis* colonies. The best explanation we have for the differences is because the CFU/mL calculation from cytometer data involves large multiplication factors. As an artefact, the calculation of just a few mislabelled “live” cells from cytometer data consequently produces falsely high estimated cell counts. For example, only one mislabelled live cell at a 100-fold dilution in the cytometer estimates 10^5^ cells/mL (1 cell/µL * 100-fold dilution * 1000µL/mL = 10^5^ cells/mL). Viewing the data on a log10 scale (Fig. S4) and inspection of the data in R reveals that this is especially a problem for *L. thermotolerans,* but also slightly for *S. cerevisiae*. Similar problems were considered and discussed by Pourcelot 2023, who concluded that cytometer counts below 10^4^ cells/mL could not be trusted (Pourcelot 2023). We therefore chose to focus only on the maximum CFU reached, which are above 10^4^ cells/mL, to minimise or eliminate the effect on our overall conclusions by incorrectly labelled “live” cells. The potential overestimation of live cells must be considered in other future experiments using fluorescent flow cytometry, especially when the bias differs between fluorescent markings (as seen here for *L. thermotolerans*).

Overall, we saw a determinable hierarchy of dominance across our four resident yeast species. Like in our communities, strongly hierarchical competitive exclusion has also been observed as a common outcome in experimental *bacterial* communities (Chang *et al*. 2023). If such competitive exclusion is found in bacterial communities with common complex metabolic interdependencies (Zelezniak *et al*. 2015; Zengler and Zaramela 2018), it makes sense that the same occurs in yeast communities; weak, or even no, niche breadth versus growth rate trade-offs in yeasts (Opulente *et al*. 2024) would predict that simply the strongest fermenting yeast species should dominate, which was seen in our experiment’s cytometer data. Our research contributes to an underrepresented yet growing research field of synthetic yeast communities (Walker and Pretorius 2022b; Pourcelot *et al*. 2023; Ruiz *et al*. 2023). Future engineering of microbial communities - yeast, bacterial, or mixed - must consider when there is a tendency for hierarchical dominance and disproportionate impacts by keystone species. If strong identity effects override the influence of species richness, it becomes logical to consider simpler synthetic communities of fewer species.

## CONCLUSIONS

We built upon previous research of species interactions in diverse wine yeast communities (Boynton and Greig 2016; Walker and Pretorius 2022a; Pourcelot *et al*. 2023, 2023; Ruiz *et al*. 2023) with the research aim to test the impact of species richness versus identity on resistance to invasion by introduced species. Using synthetic communities comprised of all combinations of four wine yeasts (*S. cerevisiae, L. thermotolerans, T. delbrueckii, S. bacillaris*), we included and tracked the presence of *B. bruxellensis* yeast and *L. plantarum* lactic acid bacteria over 21 days. In line with our predictions and previous work on the functional impact of *S. cerevisiae* (Boynton and Greig 2016), we found that species identity rather than richness drove the prevention of establishment of *B. bruxellensis* and *L. plantarum*, with *S. cerevisiae* playing a critical keystone species role. Aside from *S. cerevisiae*, our results suggest a most promising role for *L. thermotolerans* as a non-*S. cerevisiae* yeast to include in wine fermentation starter cultures. In addition to spoilage prevention by *S. cerevisiae*, the four resident yeast species demonstrated a strict dominance ranking of competitive exclusion regardless of background community composition.

A unique aspect of our study is its use of synthetic yeast communities, whereas thus far, most microbial community ecology research focuses on bacterial communities (Fiegna *et al*. 2015; Piccardi, Vessman and Mitri 2019; Hu *et al*. 2022; Chang *et al*. 2023). If the metabolic overlap across yeasts versus across bacteria differs, this shifts our predictions about the influence of species richness and niche exclusion on community resistance to invasion. Yeast species arguably consume and produce similar metabolites but just at different efficiencies (Opulente *et al*. 2024), whereas in comparison, bacteria have more unique metabolic capacities since different strains consume and produce specific metabolites, leading to cross-feeding interdependencies (Zelezniak *et al*. 2015; D’Souza *et al*. 2018; Zengler and Zaramela 2018; Kost *et al*. 2023). The association of species richness to stability and function is therefore predictably weaker in yeast communities than bacteria. Our conclusion of a keystone species rather than species richness effect on community invasion makes sense in the context of our yeast synthetic communities. We also realise that the absent effect of richness could be a result of the small number of yeast species tested here (maximum richness = 4) and that the outcome could differ if more species were included. Furthermore, most studies do not assess the entire process of invasion to a microbial community and only consider snap-shots at one or few time points (van Elsas *et al*. 2012; Mallon *et al*. 2015; Amor, Ratzke and Gore 2020). We assessed at three time points, which revealed potentially missed dynamics, such as the eventual exclusion of *L. plantarum* regardless of community species richness and identities. Our assessment of the sustained persistence or exclusion of *B. bruxellensis* and *L. plantarum* after an extended time scale of 21 days is applicable knowledge for wine production, where spoilage can often occur even after bottling (Malfeito-Ferreira and Silva 2019).

As spontaneously fermented natural wines and diverse starter cultures gain popularity (Galati *et al*. 2019; Roudil *et al*. 2020), our research emphasises a remaining importance of *S. cerevisiae* during fermentation for resilience against *B. bruxellensis* spoilage. However, if new non-*S. cerevisiae* yeasts are also desired for their aromatic or sensory contributions, it is desirable to better promote their persistence. Here, we inoculated all species concurrently and at equal abundances, which is not realistic to real wine fermentations where *S. cerevisiae* is initially at very low abundances in natural grape must (Albergaria and Arneborg 2016; Conacher *et al*. 2021; Drumonde-Neves *et al*. 2021). It would be interesting to further test varied initial abundance ratios of *S. cerevisiae* to the other three yeasts and see the impact on both resistance against *B. bruxellensis* and desired metabolic characteristics, including those produced by lactic acid bacteria such as *L. plantarum*.

Our work contributes to a growing movement of using fermented foods as a model system in ecology and evolutionary biology (Wolfe and Dutton 2015; Alekseeva *et al*. 2021; Conacher *et al*. 2021). Community microbial ecology research is being done in diverse natural and synthetic communities with prospects for long term evolution of multispecies communities (Fiegna *et al*. 2015; Piccardi, Vessman and Mitri 2019; Hu *et al*. 2022; Chang *et al*. 2023). An exciting example in fermented foods is the evolution of wine yeast with bacteria for over 100 generations, where the emergence of mutualistic benefits was observed (du Toit *et al*. 2020). Future research could work towards promoting coexistence in our wine yeast model system, such as propagating at 24 hours before the dominance of particular species, lowering the temperature, oxygenation, or reducing SO_2_ to promote less competitive species compared to *S. cerevisiae* (Jolly, Varela and Pretorius 2014; Alonso-del-Real *et al*. 2017; Bagheri *et al*. 2020). Further tests would also need to test the stability of our unique fluorescent tags over multiple generations. Experimental evolution and tracing of species sorting in synthetic yeast communities is an especially exciting prospect since they are underrepresented compared to synthetic bacterial communities.

Our research lends evidence against the commonly predicted positive relationship between species diversity and resistance to invasion. We instead observed a strong identity effect by keystone species, which could be predicted by consideration of high resource overlap in our synthetic yeast community. We recognise that the effect of species identity is not independent from species richness, since increasing richness brings greater probability that a keystone species is included. Despite the entanglement of species richness and identity effects, we feel our conclusions demonstrate the importance to test ecological theories across varied model systems and to consider both species identity and species richness when engineering microbial communities.

## AUTHOR CONTRIBUTIONS

Experimental design and conceptual ideas were developed by AML, EP, DS, TN. Experiments and data collection were performed by AML and SG, with assistance by EP, DS, and TN. Data analysis was completed by AML and TD. Original manuscript draft written by AML, with editing by AML, EP, DS, and TN.

## ACKNOWLEDGEMENTS

We thank Diego Segond for his help with data collection and cytometry in the final replicate fermentation round.

## DATA AVAILABILITY

Source data files and codes available on GitHub (amleale/2023_montpellier): https://github.com/amleale/2023_montpellier.

## SUPPLEMENTARY (tables & figures)

**Figure S1:**
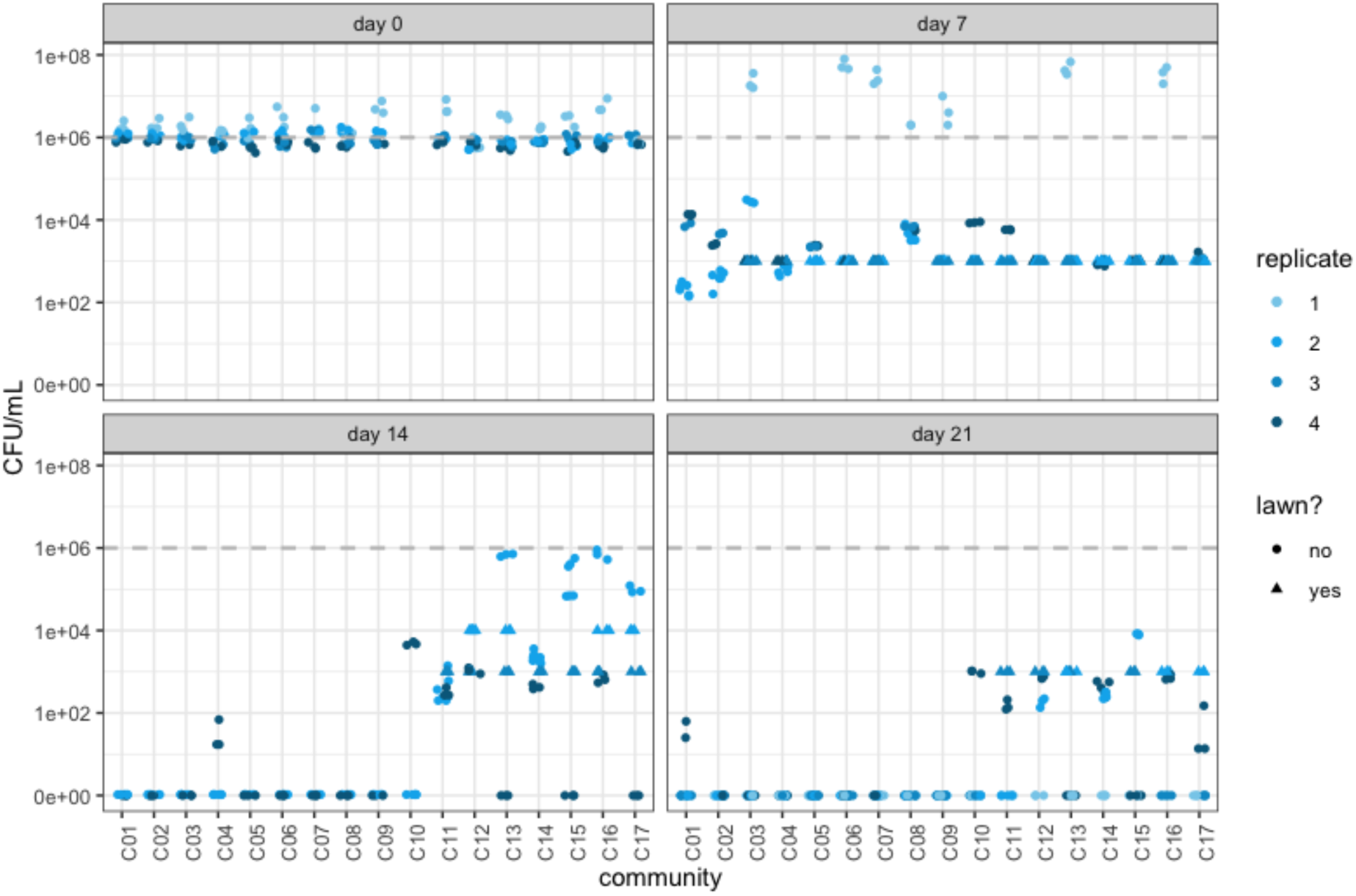
*viable yeasts remain in S. cerevisiae-free communities.* Estimated cell density of total viable yeast (excluding *B. bruxellensis*) over time from YEPD agar plating and counting after 48 hours. Dashed grey line indicates projected initial inoculated cell density at day 0 (i.e., 10^6^ cells/mL). Lawns were conservatively estimated as the minimum possible colonies (i.e., 100) and the dilution factor (i.e., a lawn at 10^-1^ dilution = 10^4^ CFU).

**Figure S2:**
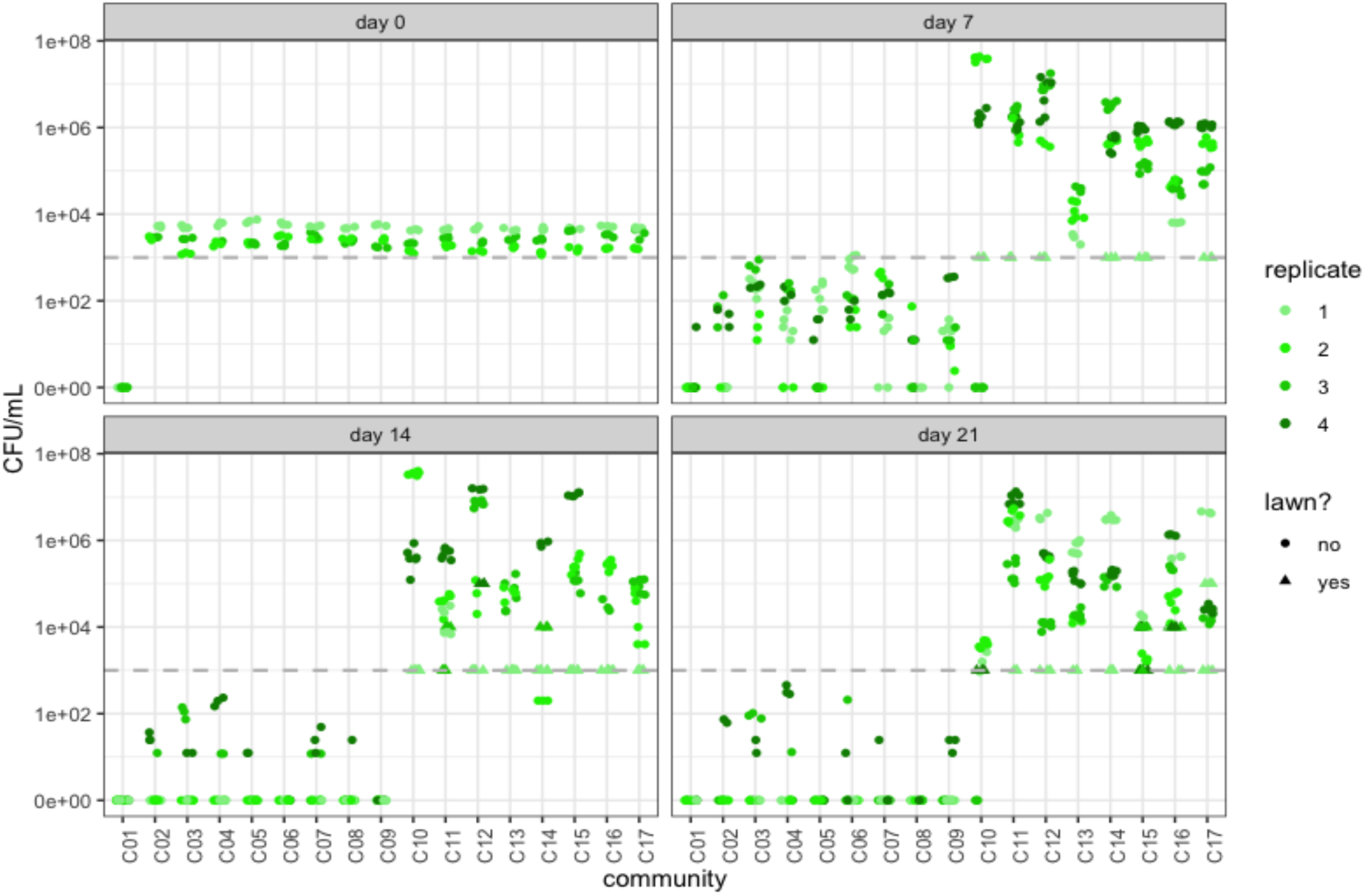
*B. bruxellensis establishes in the absence of S. cerevisiae.* Estimated cell density of *B. bruxellensis* over time from selective agar plating on YEPD supplemented with cycloheximide and counted after approximately 14 days. Dashed grey line indicates projected initial inoculated cell density of at day 0 (i.e., 10^3^ cells/mL). Lawns were conservatively estimated as the minimum possible colonies (i.e., 100) and the dilution factor (i.e., a lawn at 10^-1^ = 10^4^ CFU).

**Figure S3:**
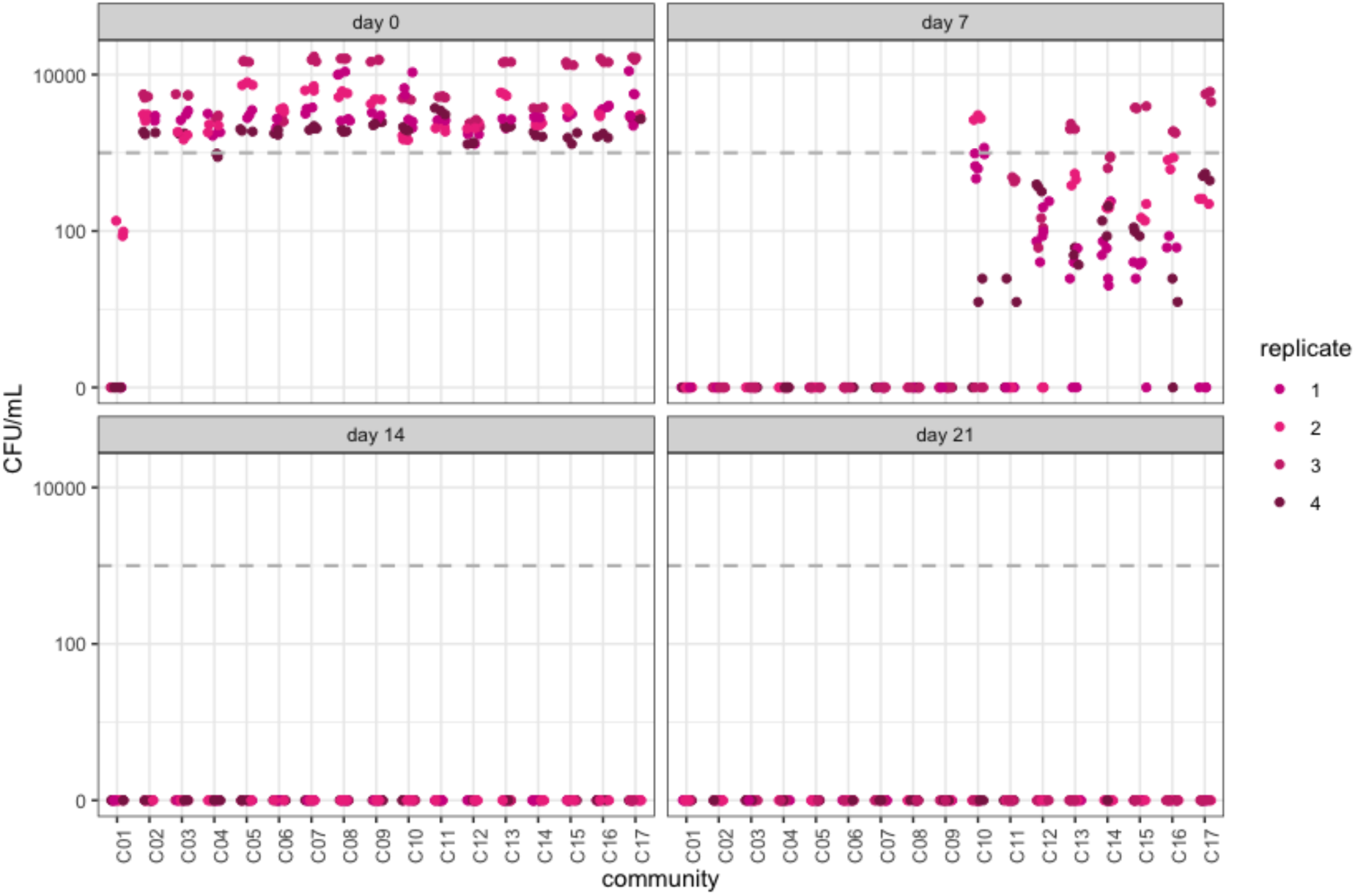
*elimination of L. plantarum is delayed in the absence of S. cerevisiae.* Estimated cell density of *L. plantarum* over time from selective agar plating on MRS supplemented with chlorophenicol and counted after 48 hours. Dashed grey line indicates projected initial inoculated cell density of at day 0 (i.e., 10^3^ cells/mL). Estimated CFU of lawns were entered depending on the dilution factor used.

**Figure S4:**
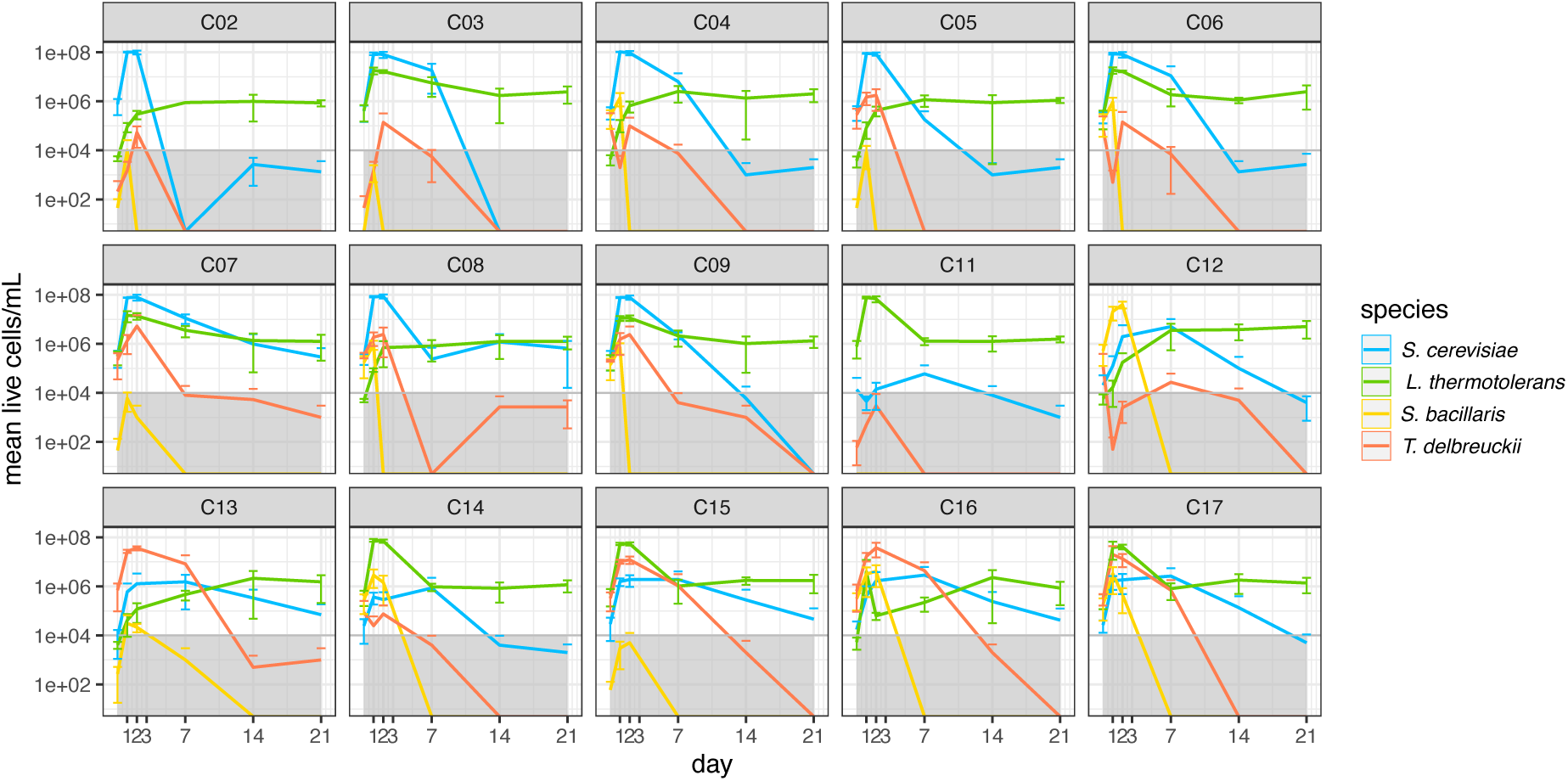
viewing main text’s Fig. 3 on log10 scale reveals biased heightened live cell counts at days 14 and 21, especially for *L. thermotolerans* and *S. cerevisiae*. Fluorescent flow cytometry estimates of resident community yeast species overtime. The mean live cells/mL of four replicate rounds of fermentation is plotted, with error bars showing one standard deviation. Grey area indicates below 10^4^ cells/mL, our proposed minimum reliable count (i.e., values below are unreliable).

**Figure S5:**
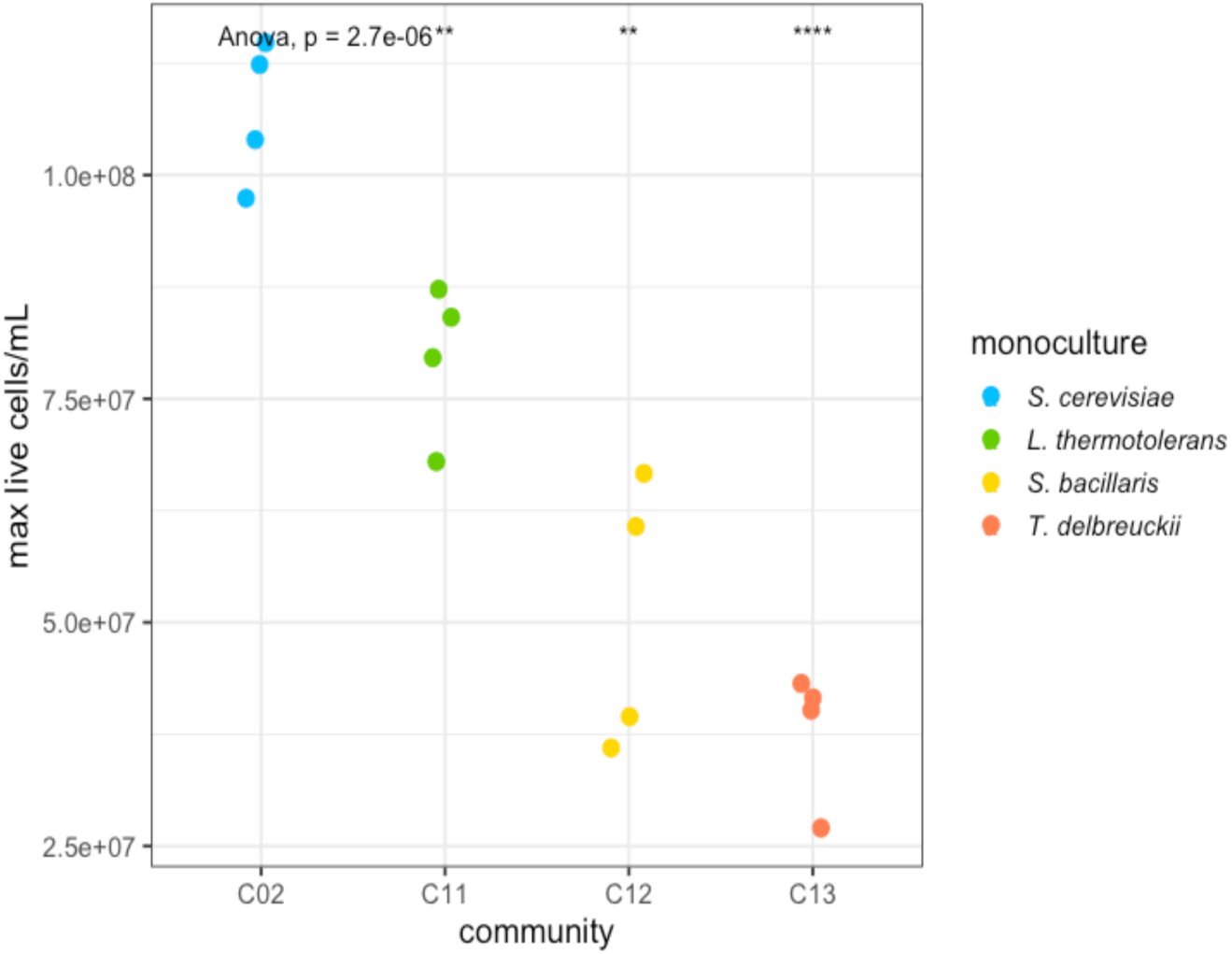
*L. thermotolerans’ monoculture reaches the greatest maximum CFU/mL of the non-S. cerevisiae species.* Fluorescent flow cytometry estimates of maximum live cells/mL reached during fermentation for the monoculture of each resident yeast species. Replication = 4 fermentation rounds per community.

**Figure S6:**
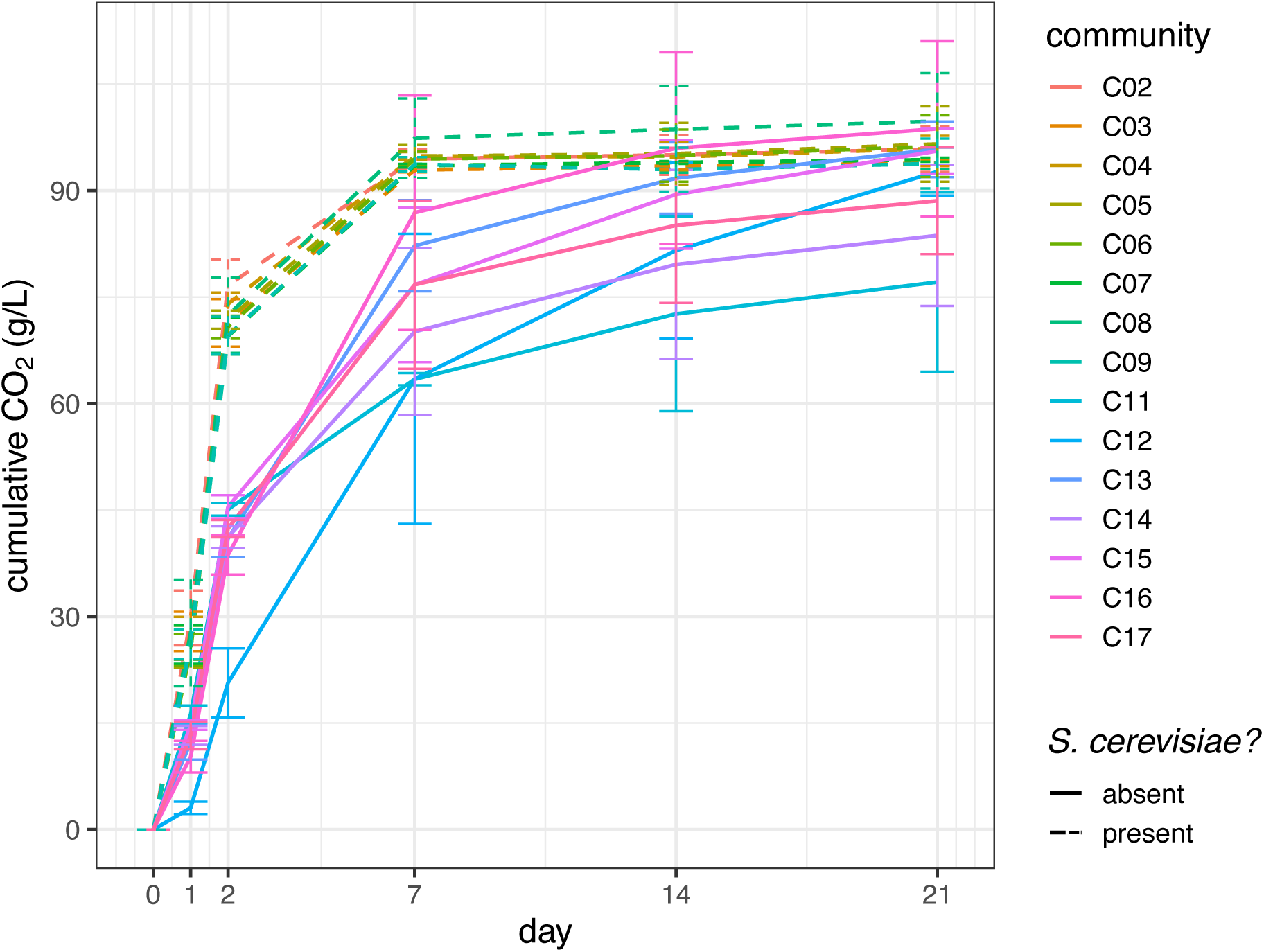
*fermentation is stunted in the absence of S. cerevisiae.* Cumulative CO_2_ produced over time demonstrates fermentation progression in all 17 synthetic communities. Each line is the mean of four replicates for each community, with error bars showing one standard deviation. Dashed lines indicate communities containing *S. cerevisiae*, solid lines are *S. cerevisiae*-free communities.

**Table S1:** p-value of Fisher’s Exact tests for all communities of detected colony growth on agar media (i.e., confounded effect of richness and *S. cerevisiae* presence). Tests completed separately for each output, day, and variable combination. Bonferroni corrected p-value significance threshold = 0.0017 (i.e., 0.05 / 30 tests). Example analysis for day 14, total viable yeast, *S. bacillaris* shown below in “Supplementary (statistical analyses), 2. Binomial plate count data”.

|  | Day | richness | *S. cerevisiae* | *L. thermotolerans* | *T. delbrueckii* | *S. bacillaris* |
| --- | --- | --- | --- | --- | --- | --- |
| Viable yeast | d07 | n/a | n/a | n/a | n/a | n/a |
|  | d14 | 0.073 | 0.004 | 0.546 | 0.044 | 0.106 |
|  | d21 | 0.001 | < 2.2e-16 | 0.736 | 0.616 | 0.177 |
| *B. bruxellensis* | d07 | 0.774 | 1.407e-09 | 2.642e-06 | 0.392 | 0.196 |
|  | d14 | 1.05e-14 | < 2.2e-16 | 0.274 | 1.00 | 0.398 |
|  | d21 | 3.425e-12 | < 2.2e-16 | 0.386 | 1.00 | 0.385 |
| *L. plantarum* | d07 | 2.709e-06 | < 2.2e-16 | 0.883 | 0.185 | 0.140 |
|  | d14 | n/a | n/a | n/a | n/a | n/a |
|  | d21 | n/a | n/a | n/a | n/a | n/a |

**Table S2:**
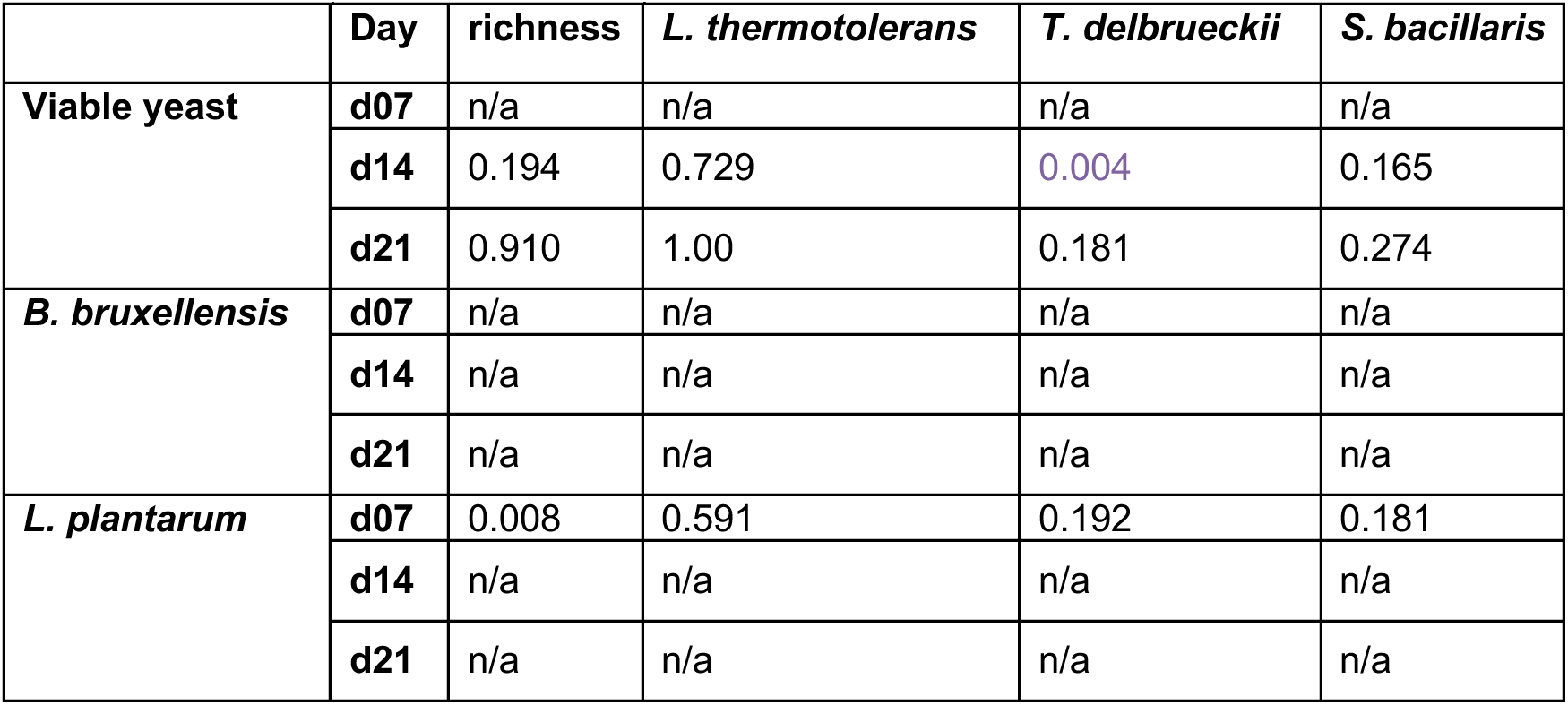
Fisher’s Exact tests for *S. cerevisiae*-free communities only (C11-C17). Tests completed separately for each output, day, and variable combination. Bonferroni corrected p-value significance threshold = 0.0042 (i.e., 0.05 / 12 tests).

**Table S3:** Fisher’s Exact tests for *S. cerevisiae*-containing communities only (C02-C09). Tests completed separately for each output, day, and variable combination. Bonferroni corrected p-value significance threshold = 0.0031 (i.e., 0.05 / 16 tests).

|  | Day | richness | *L. thermotolerans* | *T. delbrueckii* | *S. bacillaris* |
| --- | --- | --- | --- | --- | --- |
| **Viable yeast** | **d07** | n/a | n/a | n/a | n/a |
|  | **d14** | 0.825 | 0.605 | 0.605 | 0.605 |
|  | **d21** | n/a | n/a | n/a | n/a |
| ***B. bruxellensis*** | **d07** | 0.065 | 4.851e-06 | 0.1698 | 0.4923 |
|  | **d14** | 0.067 | 0.817 | 0.247 | 0.035 |
|  | **d21** | 0.231 | 0.295 | 0.034 | 1.00 |
| ***L. plantarum*** | **d07** | n/a | n/a | n/a | n/a |
|  | **d14** | n/a | n/a | n/a | n/a |
|  | **d21** | n/a | n/a | n/a | n/a |

## SUPPLEMENTARY (statistical analyses)

### 1. Cumulative CO_2_

Control communities C01 and C10 were excluded throughout. We did not have sufficient replication and degrees of freedom to test species richness 4.

<u>Day 2:</u>

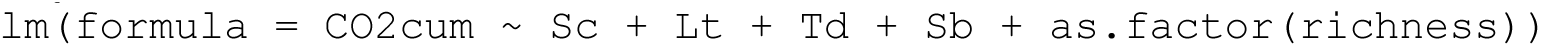

**Table S4:** Output of linear model analysis of cumulative CO2 at day 02.

|  | Estimate | SE | t-value | p-value |
| --- | --- | --- | --- | --- |
| **intercept** | 37.937 | 2.152 | 17.626 | <2e-16 |
| ***S. cerevisiae*** | 32.449 | 1.650 | 19.669 | <2e-16 |
| ***L. thermotolerans*** | 2.241 | 1.650 | 1.359 | 0.1800 |
| ***T. delbrueckii*** | 1.115 | 1.650 | 0.676 | 0.5022 |
| ***S. bacillaris*** | -4.150 | 1.650 | -2.516 | 0.0149 |
| **Richness 2** | 3.328 | 1.801 | 1.848 | 0.0702 |
| **Richness 3** | 2.106 | 2.454 | 0.858 | 0.3947 |
| **Richness 4** | NA | NA | NA | NA |

<u>Day 21:</u>

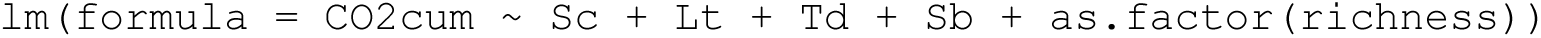

**Table S5:** Output of linear model analysis of cumulative CO_2_ at day 21.

|  | Estimate | SE | t-value | p-value |
| --- | --- | --- | --- | --- |
| **intercept** | 89.2160 | 2.5601 | 34.848 | < 2e-16 |
| ***S. cerevisiae*** | 5.1128 | 1.9623 | 2.606 | 0.01188 |
| ***L. thermotolerans*** | -5.6880 | 1.9623 | -2.899 | 0.00544 |
| ***T. delbrueckii*** | 4.3086 | 1.9623 | 2.196 | 0.03251 |
| ***S. bacillaris*** | 0.8643 | 1.9623 | 0.440 | 0.66140 |
| **Richness 2** | 2.5382 | 2.1420 | 1.185 | 0.24131 |
| **Richness 3** | 2.0704 | 2.9190 | 0.709 | 0.48126 |
| **Richness 4** | NA | NA | NA | NA |

<u>Coefficient of variation:</u>

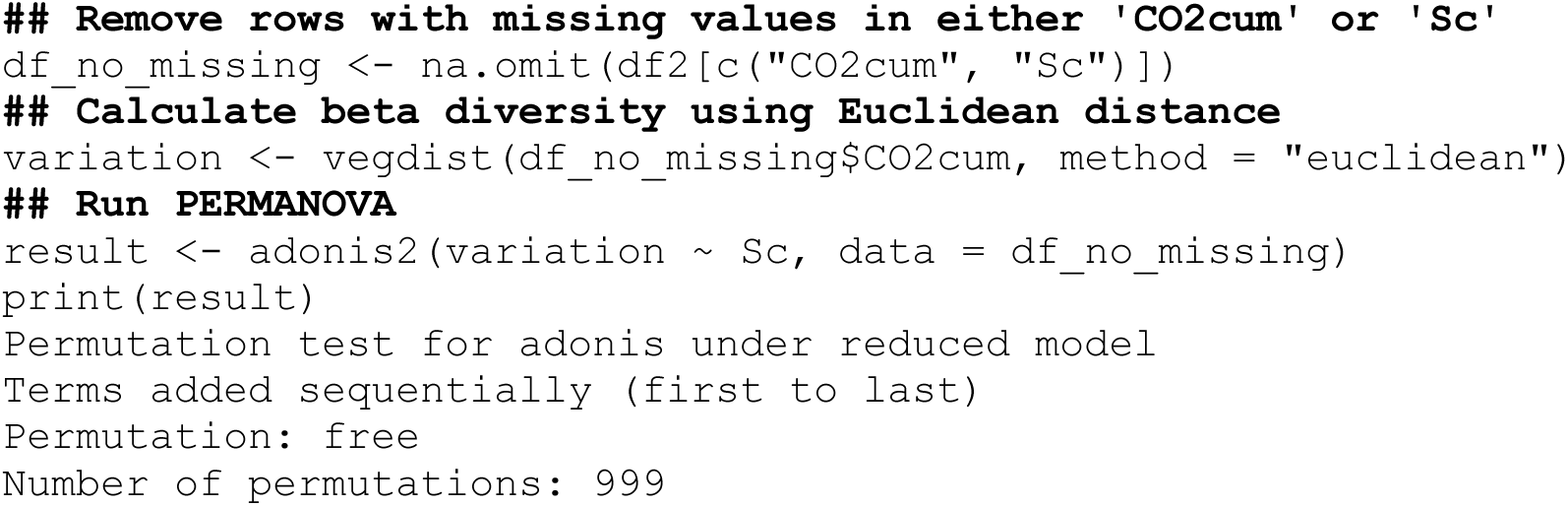

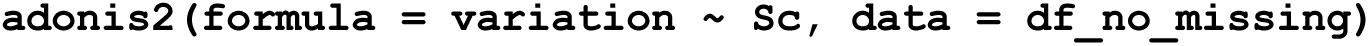

**Table S6:** Output of PERMANOVA test demonstrates variation in cumulative CO_2_ differs between *S. cerevisiae*-free versus *S. cerevisiae*-containing communities.

|  | DF | SumSq | R2 | F | p-value |
| --- | --- | --- | --- | --- | --- |
| ***S. cerevisiae*** | 1 | 17299 | 0.03364 | 12.464 | 0.002 |
| **Residual** | 358 | 496879 | 0.96636 |  |  |

<u>All time points:</u>

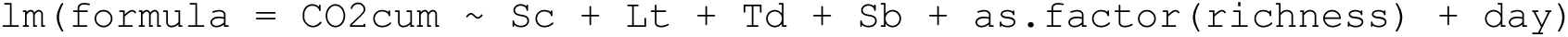

**Table S7:** Output of linear model analysis of cumulative CO_2_ for all communities and time points.

|  | Estimate | SE | t-value | p-value |
| --- | --- | --- | --- | --- |
| **intercept** | 19.6969 | 3.6370 | 5.416 | 1.13e-07 |
| ***S. cerevisiae*** | 13.6592 | 2.6405 | 5.173 | **3.88e-07** |
| ***L. thermotolerans*** | -1.6775 | 2.6405 | -0.635 | 0.526 |
| ***T. delbrueckii*** | 2.9111 | 2.6405 | 1.102 | 0.271 |
| ***S. bacillaris*** | -0.7964 | 2.6405 | -0.302 | 0.763 |
| **Richness 2** | 2.4290 | 2.8823 | 0.843 | 0.400 |
| **Richness 3** | 1.7110 | 3.9279 | 0.436 | 0.663 |
| **Richness 4** | NA | NA | NA | NA |
| **day** | 3.8529 | 0.1555 | 24.781 | **< 2e-16** |

<u>Day 07 - *S. cerevisiae-*containing communities only:</u>

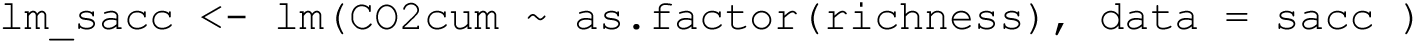

**Table S8:** Output of linear model analysis of cumulative CO_2_ at day 07 for *S. cerevisiae*-containing communities only (C02-C09).

|  | **Estimate** | **SE** | **t-value** | **p-value** |
| --- | --- | --- | --- | --- |
| **Intercept** | 94.4639 | 1.1887 | 79.470 | <2e-16 |
| **Richness 2** | -0.2490 | 1.3726 | -0.181 | 0.857 |
| **Richness 3** | 0.6506 | 1.3726 | 0.474 | 0.639 |
| **Richness 4** | -0.7602 | 1.6810 | -0.452 | 0.655 |

### 2. Binomial plate count data

used to create supplementary Tables S1, S2, S3

<u>Example of Fisher’s Test:</u> (day 14, total viable yeast, *S. bacillaris*)

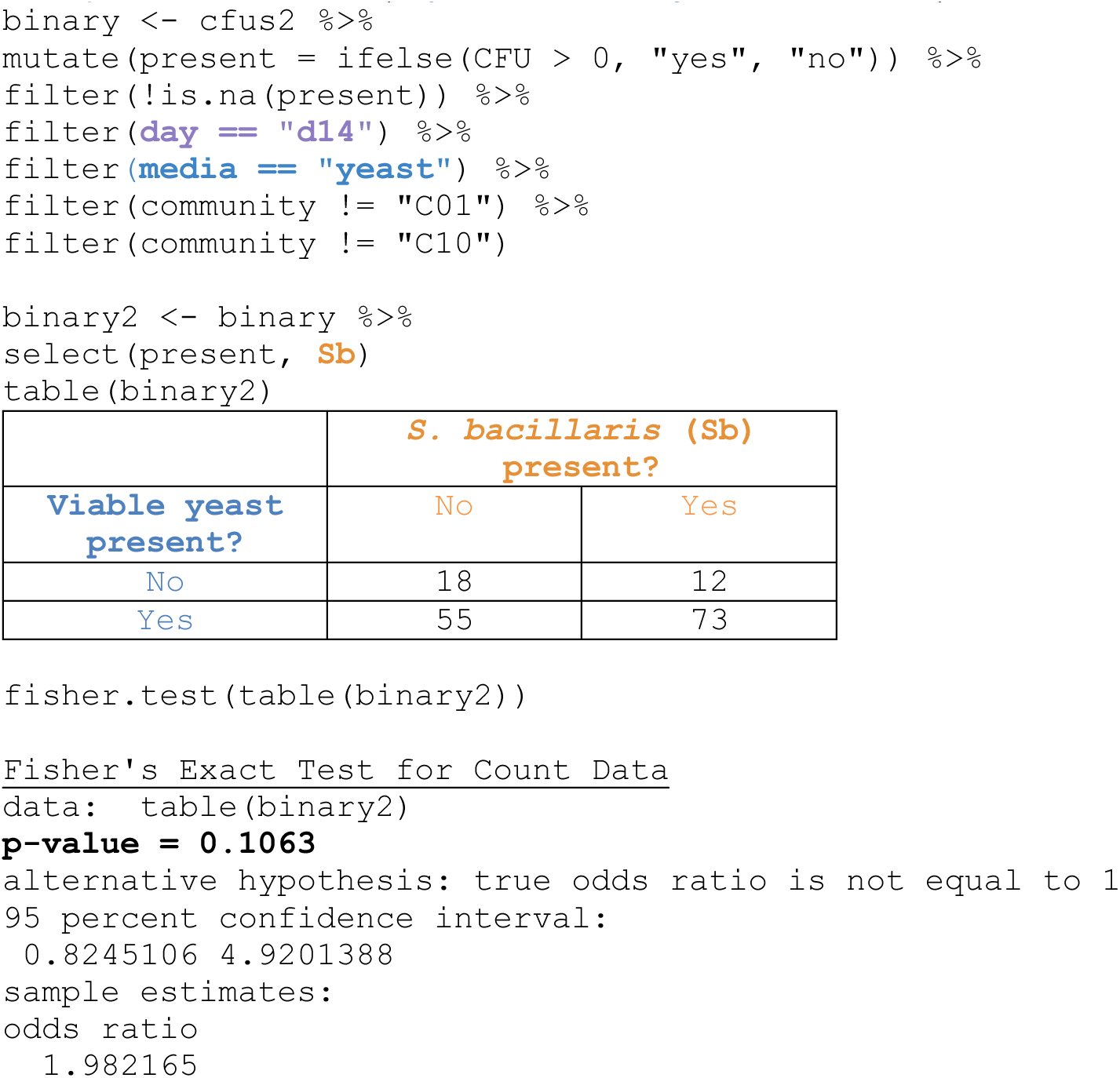

## REFERENCES

Albergaria H, Arneborg N. Dominance of Saccharomyces cerevisiae in alcoholic fermentation processes: role of physiological fitness and microbial interactions. Appl Microbiol Biotechnol 2016;100:2035–46.

Alekseeva AY, Groenenboom AE, Smid EJ et al. Eco-Evolutionary Dynamics in Microbial Communities from Spontaneous Fermented Foods. Int J Environ Res Public Health 2021;18:10093.

Allison SD, Martiny JBH. Resistance, resilience, and redundancy in microbial communities. Proc Natl Acad Sci 2008;105:11512–9.

Alonso-del-Real J, Lairón-Peris M, Barrio E et al. Effect of Temperature on the Prevalence of Saccharomyces Non cerevisiae Species against a S. cerevisiae Wine Strain in Wine Fermentation: Competition, Physiological Fitness, and Influence in Final Wine Composition. Front Microbiol 2017;8.

Amor DR, Ratzke C, Gore J. Transient invaders can induce shifts between alternative stable states of microbial communities. Sci Adv 2020;6:eaay8676.

Bagheri B, Bauer FF, Cardinali G et al. Ecological interactions are a primary driver of population dynamics in wine yeast microbiota during fermentation. Sci Rep 2020;10:4911.

Bartowsky E j., Costello P j., Chambers P j. Emerging trends in the application of malolactic fermentation. Aust J Grape Wine Res 2015;21:663–9.

Bates D, Mächler M, Bolker B et al. Fitting Linear Mixed-Effects Models Using lme4. J Stat Softw 2015;67:1–48.

Berry D, Widder S. Deciphering microbial interactions and detecting keystone species with co-occurrence networks. Front Microbiol 2014;5.

Binati RL, Lemos Junior WJF, Luzzini G et al. Contribution of non-Saccharomyces yeasts to wine volatile and sensory diversity: A study on Lachancea thermotolerans, Metschnikowia spp. and Starmerella bacillaris strains isolated in Italy. Int J Food Microbiol 2020;318:108470.

Boynton PJ, Greig D. Species richness influences wine ecosystem function through a dominant species. Fungal Ecol 2016;22:61–72.

Bravo-Ferrada BM, Hollmann A, Delfederico L et al. Patagonian red wines: selection of Lactobacillus plantarum isolates as potential starter cultures for malolactic fermentation. World J Microbiol Biotechnol 2013;29:1537–49.

Cappello MS, Zapparoli G, Logrieco A et al. Linking wine lactic acid bacteria diversity with wine aroma and flavour. Int J Food Microbiol 2017;243:16–27.

Chang C-Y, Bajić D, Vila JCC et al. Emergent coexistence in multispecies microbial communities. Science 2023;381:343–8.

Childs BC, Bohlscheid JC, Edwards CG. Impact of available nitrogen and sugar concentration in musts on alcoholic fermentation and subsequent wine spoilage by *Brettanomyces bruxellensis*. Food Microbiol 2015;46:604–9.

Ciani M, Beco L, Comitini F. Fermentation behaviour and metabolic interactions of multistarter wine yeast fermentations. Int J Food Microbiol 2006;108:239–45.

Ciani M, Capece A, Comitini F et al. Yeast Interactions in Inoculated Wine Fermentation. Front Microbiol 2016;7.

Ciani M, Comitini F, Mannazzu I et al. Controlled mixed culture fermentation: a new perspective on the use of non-Saccharomyces yeasts in winemaking. FEMS Yeast Res 2010;10:123–33.

Conacher C, Luyt N, Naidoo-Blassoples R et al. The ecology of wine fermentation: a model for the study of complex microbial ecosystems. Appl Microbiol Biotechnol 2021;105:3027–43.

Csoma H, Sipiczki M. Taxonomic reclassification of Candida stellata strains reveals frequent occurrence of Candida zemplinina in wine fermentation. FEMS Yeast Res 2008;8:328–36.

D’Antonio CM, Thomsen M. Ecological Resistance in Theory and Practice. Weed Technol 2004;18:1572–7.

Donohue I, Hillebrand H, Montoya JM et al. Navigating the complexity of ecological stability. Ecol Lett 2016;19:1172–85.

Drumonde-Neves J, Fernandes T, Lima T et al. Learning from 80 years of studies: a comprehensive catalogue of non-Saccharomyces yeasts associated with viticulture and winemaking. FEMS Yeast Res 2021;21:foab017.

D’Souza G, Shitut S, Preussger D et al. Ecology and evolution of metabolic cross-feeding interactions in bacteria. Nat Prod Rep 2018;35:455–88.

Du Toit M, Pretorius IS. Microbial spoilage and preservation of wine : using weapons for nature’s own arsenal. 2000.

Eisenhauer N, Schulz W, Scheu S et al. Niche dimensionality links biodiversity and invasibility of microbial communities. Funct Ecol 2013;**27**:282–8.

van Elsas JD, Chiurazzi M, Mallon CA et al. Microbial diversity determines the invasion of soil by a bacterial pathogen. Proc Natl Acad Sci 2012;109:1159–64.

Elton CS. The Ecology of Invasions by Animals and Plants. Boston, MA: Springer US, 1958.

Emery SM, Gross KL. Dominant Species Identity, Not Community Evenness, Regulates Invasion in Experimental Grassland Plant Communities. Ecology 2007;**88**:954–64.

Englezos V, Cocolin L, Rantsiou K et al. Specific Phenotypic Traits of Starmerella bacillaris Related to Nitrogen Source Consumption and Central Carbon Metabolite Production during Wine Fermentation. Appl Environ Microbiol 2018;84:e00797–18.

Englezos V, Jolly NP, Di Gianvito P et al. Microbial interactions in winemaking: Ecological aspects and effect on wine quality. Trends Food Sci Technol 2022;127:99–113.

Ernst AR, Barak RS, Glasenhardt M-C et al. Dominant species establishment may influence invasion resistance more than phylogenetic or functional diversity. J Appl Ecol 2023;60:2652–64.

Ernst AR, Barak RS, Hipp AL et al. The invasion paradox dissolves when using phylogenetic and temporal perspectives. J Ecol 2022;110:443–56.

Fiegna F, Moreno-Letelier A, Bell T et al. Evolution of species interactions determines microbial community productivity in new environments. ISME J 2015;9:1235–45.

Fleet GH. Yeast interactions and wine flavour. Int J Food Microbiol 2003;86:11–22.

Fridley JD, Stachowicz JJ, Naeem S et al. The Invasion Paradox: Reconciling Pattern and Process in Species Invasions. Ecology 2007;88:3–17.

Galati A, Schifani G, Crescimanno M et al. “Natural wine” consumers and interest in label information: An analysis of willingness to pay in a new Italian wine market segment. J Clean Prod 2019;227:405–13.

G-Alegría E, López I, Ruiz JI et al. High tolerance of wild *Lactobacillus plantarum* and *Oenococcus oeni* strains to lyophilisation and stress environmental conditions of acid pH and ethanol. FEMS Microbiol Lett 2004;230:53–61.

Harrouard J, Eberlein C, Ballestra P et al. Brettanomyces bruxellensis: Overview of the genetic and phenotypic diversity of an anthropized yeast. Mol Ecol 2023;32:2374–95.

Hu J, Amor DR, Barbier M et al. Emergent phases of ecological diversity and dynamics mapped in microcosms. Science 2022;378:85–9.

Ivey M, Massel M, Phister TG. Microbial Interactions in Food Fermentations. Annu Rev Food Sci Technol 2013;4:141–62.

Jiang L, Morin PJ. Productivity gradients cause positive diversity–invasibility relationships in microbial communities. Ecol Lett 2004;7:1047–57.

Jolly NP, Varela C, Pretorius IS. Not your ordinary yeast: non-Saccharomyces yeasts in wine production uncovered. FEMS Yeast Res 2014;14:215–37.

Jousset A, Schmid B, Scheu S et al. Genotypic richness and dissimilarity opposingly affect ecosystem functioning. Ecol Lett 2011;14:537–45.

Kemsawasd V, Viana T, Ardö Y et al. Influence of nitrogen sources on growth and fermentation performance of different wine yeast species during alcoholic fermentation. Appl Microbiol Biotechnol 2015;99:10191–207.

Kennedy TA, Naeem S, Howe KM et al. Biodiversity as a barrier to ecological invasion. Nature 2002;417:636–8.

Kinnunen M, Dechesne A, Proctor C et al. A conceptual framework for invasion in microbial communities. ISME J 2016;10:2773–9.

Kost C, Patil KR, Friedman J et al. Metabolic exchanges are ubiquitous in natural microbial communities. Nat Microbiol 2023;8:2244–52.

Liu J, Huang T-Y, Liu G et al. Microbial Interaction between Lactiplantibacillus plantarum and Saccharomyces cerevisiae: Transcriptome Level Mechanism of Cell-Cell Antagonism. Microbiol Spectr 2022;10:e01433–22.

Lonvaud-Funel A. Lactic acid bacteria in the quality improvement and depreciation of wine. In: Konings WN, Kuipers OP, In ’t Veld JHJH (eds.). Lactic Acid Bacteria: Genetics, Metabolism and Applications: Proceedings of the Sixth Symposium on Lactic Acid Bacteria: Genetics, Metabolism and Applications, 19–23 September 1999, Veldhoven, The Netherlands. Dordrecht: Springer Netherlands, 1999, 317–31.

Malfeito-Ferreira M, Silva AC. Spoilage Yeasts in Wine Production. In: Romano P, Ciani M, Fleet GH (eds.). Yeasts in the Production of Wine. New York, NY: Springer, 2019, 375–94.

Mallon CA, Elsas JDV, Salles JF. Microbial Invasions: The Process, Patterns, and Mechanisms. Trends Microbiol 2015;23:719–29.

Mallon CA, Poly F, Le Roux X et al. Resource pulses can alleviate the biodiversity–invasion relationship in soil microbial communities. Ecology 2015;96:915–26.

Medina K, Boido E, Dellacassa E et al. Growth of non-Saccharomyces yeasts affects nutrient availability for Saccharomyces cerevisiae during wine fermentation. Int J Food Microbiol 2012;157:245–50.

Mills LS, Soulé ME, Doak DF. The Keystone-Species Concept in Ecology and Conservation. BioScience 1993;43:219–24.

Morneau AD, Zuehlke JM, Edwards CG. Comparison of media formulations used to selectively cultivate Dekkera/Brettanomyces. Lett Appl Microbiol 2011;53:460–5.

Mouquet N, Gravel D, Massol F et al. Extending the concept of keystone species to communities and ecosystems. Ecol Lett 2013;16:1–8.

Ngwenya MP, Nkambule TP, Kidane SW. Physicochemical attributes and acceptability of marula wine fermented with natural *Lactiplantibacillus plantarum* and *Saccharomyces cerevisiae*. Heliyon 2023;9:e21613.

Opulente DA, LaBella AL, Harrison M-C et al. Genomic factors shape carbon and nitrogen metabolic niche breadth across Saccharomycotina yeasts. Science 2024;384:eadj4503.

Pereira PR, Freitas CS, Paschoalin VMF. Saccharomyces cerevisiae biomass as a source of next-generation food preservatives: Evaluating potential proteins as a source of antimicrobial peptides. Compr Rev Food Sci Food Saf 2021;20:4450–79.

Petruzzella A, da S. S. R. Rodrigues TA, van Leeuwen CHA et al. Species identity and diversity effects on invasion resistance of tropical freshwater plant communities. Sci Rep 2020;10:5626.

Philippot L, Griffiths BS, Langenheder S. Microbial Community Resilience across Ecosystems and Multiple Disturbances. Microbiol Mol Biol Rev 2021;**85**:10.1128/mmbr.00026-20.

Piccardi P, Vessman B, Mitri S. Toxicity drives facilitation between 4 bacterial species. Proc Natl Acad Sci 2019;116:15979–84.

Pourcelot E. Développement d’une communauté modèle de levures oenologiques. 2023.

Pourcelot E, Conacher C, Marlin T et al. Comparing the hierarchy of inter- and intra-species interactions with population dynamics of wine yeast cocultures. FEMS Yeast Res 2023;23:foad039.

Pourcelot E, Vigna A, Marlin T et al. Design of a new model yeast consortium for ecological studies of enological fermentation. 2024:2024.05.06.592697.

Ramírez M, Velázquez R. The Yeast Torulaspora delbrueckii: An Interesting But Difficult-To-Use Tool for Winemaking. Fermentation 2018;4:94.

Ratzke C, Barrere J, Gore J. Strength of species interactions determines biodiversity and stability in microbial communities. Nat Ecol Evol 2020;4:376–83.

Renouf V, Falcou M, Miot-Sertier C et al. Interactions between Brettanomyces bruxellensis and other yeast species during the initial stages of winemaking. J Appl Microbiol 2006;100:1208–19.

Roudil L, Russo P, Berbegal C et al. Non-Saccharomyces Commercial Starter Cultures: Scientific Trends, Recent Patents and Innovation in the Wine Sector. 2020, DOI: 10.2174/2212798410666190131103713.

van Ruijven J, De Deyn GB, Berendse F. Diversity reduces invasibility in experimental plant communities: the role of plant species. Ecol Lett 2003;6:910–8.

Ruiz J, de Celis M, Diaz-Colunga J et al. Predictability of the community-function landscape in wine yeast ecosystems. Mol Syst Biol 2023;19:e11613.

Shekhawat K, Bauer FF, Setati ME. Impact of oxygenation on the performance of three non-Saccharomyces yeasts in co-fermentation with Saccharomyces cerevisiae. Appl Microbiol Biotechnol 2017;101:2479–91.

Smith BD, Divol B. *Brettanomyces bruxellensis*, a survivalist prepared for the wine apocalypse and other beverages. Food Microbiol 2016;59:161–75.

Taillandier P, Lai QP, Julien-Ortiz A et al. Interactions between Torulaspora delbrueckii and Saccharomyces cerevisiae in wine fermentation: influence of inoculation and nitrogen content. World J Microbiol Biotechnol 2014;30:1959–67.

Thébault E, Loreau M. Trophic Interactions and the Relationship between Species Diversity and Ecosystem Stability. Am Nat 2005;166:E95–114.

Tilman D. Niche tradeoffs, neutrality, and community structure: a stochastic theory of resource competition, invasion, and community assembly. Proc Natl Acad Sci 2004;101:10854–61.

du Toit M, Engelbrecht L, Lerm E et al. Lactobacillus: the Next Generation of Malolactic Fermentation Starter Cultures—an Overview. Food Bioprocess Technol 2011;4:876–906.

du Toit SC, Rossouw D, du Toit M et al. Enforced Mutualism Leads to Improved Cooperative Behavior between Saccharomyces cerevisiae and Lactobacillus plantarum. Microorganisms 2020;8:1109.

Urbina Á, Calderón F, Benito S. The Combined Use of Lachancea thermotolerans and Lactiplantibacillus plantarum (former Lactobacillus plantarum) in Wine Technology. Foods 2021;10:1356.

Urso R, Rantsiou K, Dolci P et al. Yeast biodiversity and dynamics during sweet wine production as determined by molecular methods. FEMS Yeast Res 2008;8:1053–62.

Vila JC, Jones ML, Patel M et al. Uncovering the rules of microbial community invasions. Nat Ecol Evol 2019;3:1162–71.

Virdis C, Sumby K, Bartowsky E et al. Lactic Acid Bacteria in Wine: Technological Advances and Evaluation of Their Functional Role. Front Microbiol 2021;11.

Walker RS, Pretorius IS. Synthetic biology for the engineering of complex wine yeast communities. Nat Food 2022a;3:249–54.

Walker RSK, Pretorius IS. Synthetic biology for the engineering of complex wine yeast communities. Nat Food 2022b;3:249–54.

Wedral D, Shewfelt R, Frank J. The challenge of Brettanomyces in wine. LWT - Food Sci Technol 2010;43:1474–9.

Weiss AS, Niedermeier LS, von Strempel A et al. Nutritional and host environments determine community ecology and keystone species in a synthetic gut bacterial community. Nat Commun 2023;14:4780.

Williams KM, Liu P, Fay JC. Evolution of ecological dominance of yeast species in high-sugar environments. Evolution 2015;69:2079–93.

Wolfe BE, Dutton RJ. Fermented foods as experimentally tractable microbial ecosystems. Cell 2015;161:49–55.

Xing J, Jia X, Wang H et al. The legacy of bacterial invasions on soil native communities. Environ Microbiol 2021;23:669–81.

Zelezniak A, Andrejev S, Ponomarova O et al. Metabolic dependencies drive species co-occurrence in diverse microbial communities. Proc Natl Acad Sci 2015;112:6449–54.

Zengler K, Zaramela LS. The social network of microorganisms — how auxotrophies shape complex communities. Nat Rev Microbiol 2018;16:383–90.

Zott K, Miot-Sertier C, Claisse O et al. Dynamics and diversity of non-*Saccharomyces* yeasts during the early stages in winemaking. Int J Food Microbiol 2008;125:197–203.

